# Causal Cortical and Thalamic Connections in the Human Brain

**DOI:** 10.1101/2024.06.22.600166

**Authors:** Dian Lyu, James Stiger, Zoe Lusk, Vivek Buch, Josef Parvizi

## Abstract

The brain’s functional architecture is intricately shaped by causal connections between its cortical and subcortical structures. Here, we studied 27 participants with 4864 electrodes implanted across the anterior, mediodorsal, and pulvinar thalamic regions, and the cortex. Using data from electrical stimulation procedures and a data-driven approach informed by neurophysiological standards, we dissociated three unique spectral patterns generated by the perturbation of a given brain area. Among these, a novel waveform emerged, marked by delayed-onset slow oscillations in both ipsilateral and contralateral cortices following thalamic stimulations, suggesting a mechanism by which a thalamic site can influence bilateral cortical activity. Moreover, cortical stimulations evoked earlier signals in the thalamus than in other connected cortical areas suggesting that the thalamus receives a copy of signals before they are exchanged across the cortex. Our causal connectivity data can be used to inform biologically-inspired computational models of the functional architecture of the brain.

## Introduction

The brain’s dynamics are shaped by electrophysiological interactions throughout its subregions, encompassing not only cortical but also subcortical regions. Advancements in neuroimaging have provided significant insights into the global architecture of the brain’s functional connectivity^1^. However, we know considerably less about the dynamic, fast-paced and causal electrophysiological relationships in the human brain. This knowledge gap is even more pronounced when it comes to understanding the role of subcortical areas such as the thalamus, which is known to play a key role in modulating the global dynamics of the brain^2^.

A classic method of studying causal electrophysiological relationships in the brain is by sending repeated single electrical pulses to intracranially implanted electrodes while recording the presence or absence of electrophysiological changes in all other areas of the brain where recording electrodes are present^3^. This method has been traditionally referred to as the study of cortico-cortical evoked potentials^4^ (CCEPs) since it has primarily been used to study causal electrophysiological connectivity between pairs of *cortical* regions. Similar studies of thalamocortical and corticothalamic connections have been rarely conducted since direct recordings from and stimulations in the human thalamus are extremely rare in human neuroscience research^5–7^. As a result, it remains unknown if stimulation of the thalamus will have a different, or the same, effect on other regions of the brain. Moreover, with a few exceptions^7,8^, previous studies have been largely reliant on simple univariate measures detecting large peaks or the time-to-peak in the evoked signals, and limit responses to fixed windows of interest. These traditional methods are not able to capture the complex dynamics of physiological responses generated by electrical stimulations (**Figure 1**); and they can vary significantly depending on the chosen cutoff z-scores for determining an effect, current intensity and the distance between stimulating and recording electrode contacts^9^. These problems become more significant when we lack prior data to inform our hypotheses about evoked potentials caused by human subcortical structures such as the thalamus.

**Figure 1.**
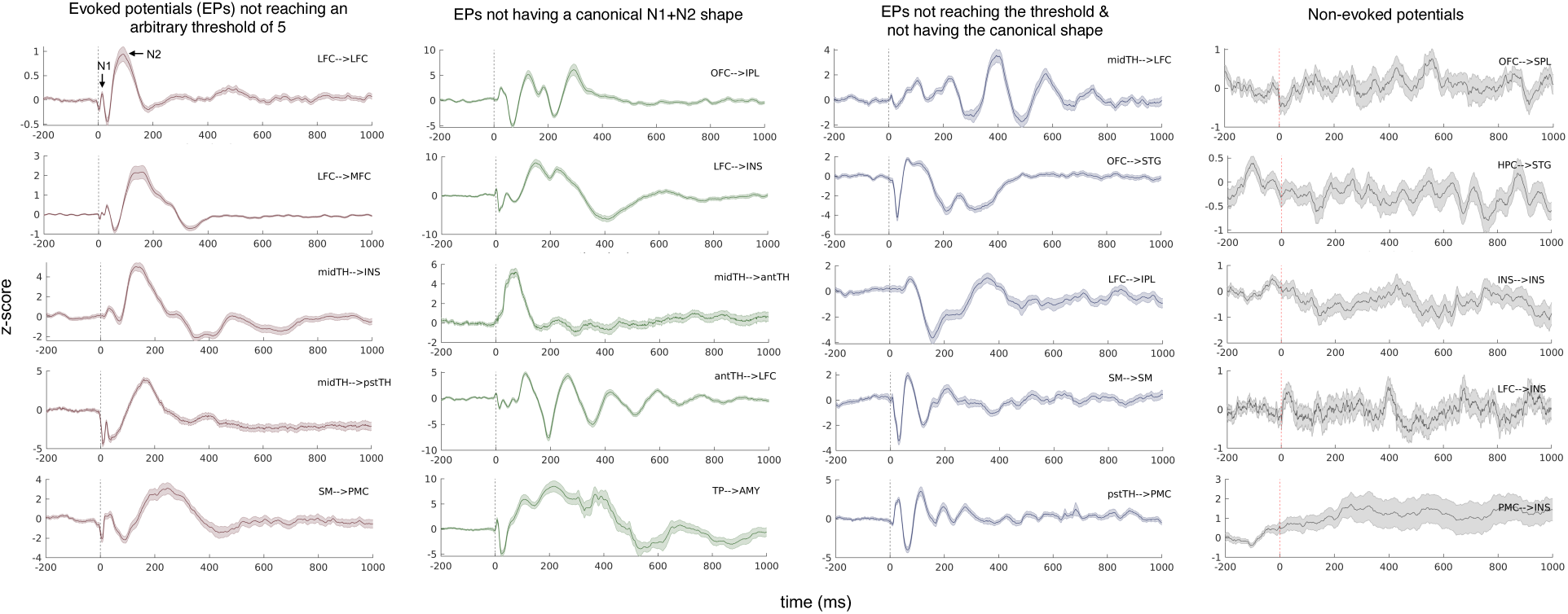
Electrophysiological responses (local field potential) evoked by the stimulation of a given brain area are complex and variable. Here we show randomly selected examples of the complex waveforms generated by electrical stimulation. A large portion of evoked responses either does not reach an arbitrary threshold, e.g., z-score = 5 that is usually used in the literature (first column, with five separate examples) or does not conform to a typical shape of evoked responses with an N1 and N2 components (second column) or neither have a large amplitude nor a canonical N1 and N2 shape (third column). By using univariate measures, one will only map a fraction of true connections in the brain. Plotted signals are the evoked responses stimulated and recorded from a pair of bipolar sites (i.e., from one stimulation bipolar site to one recording bipolar site), of which the anatomical information is provided in the upper corner. The central line is the trials-averaged signal, baseline-corrected and z-scored, while the shaded area depicts the standard error over the 45 trials.

In the current study, we aim to use a novel approach to provide a map of causal electrophysiological connections in the human brain including the thalamus.

## RESULTS

Our results are based on intracranial recordings and stimulations in twenty-seven participants (40.7% female; mean age ± SD: 34.9 ± 10.0 years) diagnosed with focal epilepsy. Each participant provided data from an average of 180 (± 46) electrode sites. The entire dataset encompasses 4864 sites across all subjects, with each adjacent pair stimulated approximately 45 times (with sufficient inter-trial intervals to avoid overlapping effects), while recordings were taken from all other available sites.

We leveraged a recently developed clinical method that involves *thalamic recordings* in neurosurgical patients undergoing invasive stereotactic electroencephalography (sEEG)^5^. In this procedure, intracranial electrodes were implanted in three different sites of the thalamus — corresponding to anterior, mediodorsal, and pulvinar nuclei — by extending electrodes initially placed in the cortical areas. This procedure was designed for extended clinical indication without increasing the number of implanted electrodes.

### Identifying Common Stable Patterns in Stimulation Evoked Potentials

To determine true physiological effects upon electrical stimulation, we used a non-linear manifold learning algorithm^10^ integrated with human judgement, based on the spectral information of evoked potentials (EPs) across time and frequencies (**Figure 2 a-c**). In this approach, we first labeled the data partially and tentatively, i.e., subject to subsequent data-driven corrections, based on two measures of connectivity: (i) trial-averaged *power* of EPs as a measure of strength, and (ii) *inter-trial phase coherence* (ITPC) across repeated stimulations as a measure of consistency for the evoked responses^11^. We applied a semi-supervised machine learning approach to decipher the inherent data structure from this preliminary labelling, and re-classified the data, within each subject, into two separate clusters (**Fig. S2**): one cluster of data where stimulation of the seed site caused significant EPs in the target site, and another cluster of data where stimulation of the seed site did not cause any significant EPs in the target site. For simplicity, we called these two clusters “*activated*” and “*non-activated”*, respectively.

**Figure 2.**
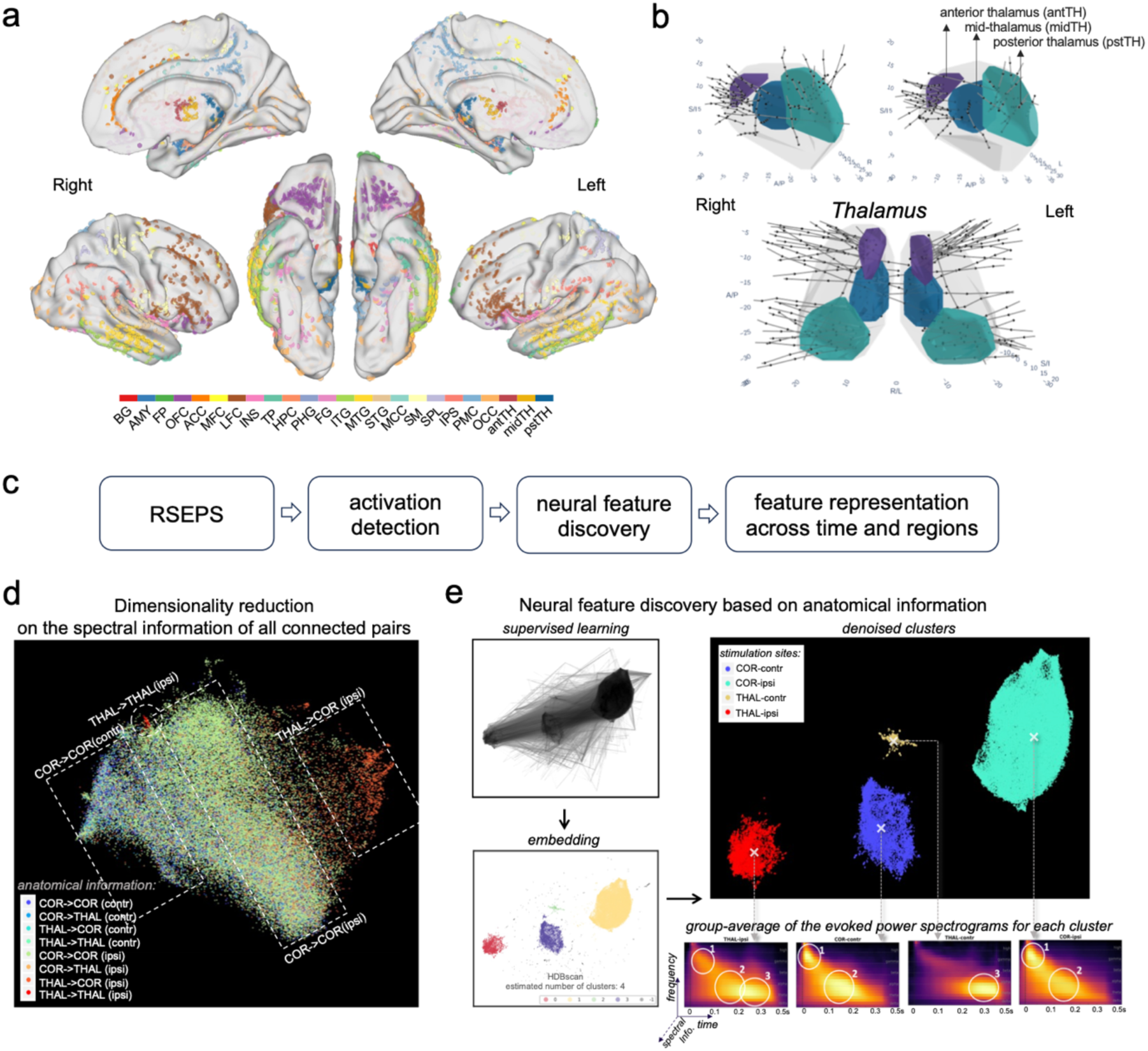
Illustration of data processing pipeline. (a) Electrode coverage at the group level. The electrode localization was manually labelled by the experienced neurologist on the team, based on each subject’s brain morphology in their own high-resolution T1 scan (L: left, R: right, ant: anterior, pst: posterior, PMC: posteromedial cortex, SM: sensorimotor, SPL: superior parietal lobule, ACC: anterior cingulate cortex, CLT: claustrum, TH: thalamus, IPL: inferior parietal lobule, MFC: medial frontal cortex, OFC: orbital frontal cortex, STG: superior temporal gyrus, LFC: lateral frontal cortex, INS: insula, FG: fusiform gyrus, HPC: hippocampus, MTG: middle temporal gyrus, PHG: parahippocampal gyrus, AMY: amygdala, TP: temporal pole, MCC: midcingulate cortex, ITG: inferior temporal gyrus). For visualization purposes, each subject’s brain images were normalized to a standard brain surface space (FS_LR: https://osf.io/k89fh/wiki/Surface/) with FreeSurfer (https://surfer.nmr.mgh.harvard.edu/) and Connectome-Workbench (https://www.humanconnectome.org/software/connectome-workbench), and their electrodes were projected to the standard surface (sites out of grey matter are excluded). **(b)** The localization of electrodes in the thalamus, divided into three thalamic divisions: anterior (purple), mid (blue), and posterior (green) subregions of the thalamus. **(c)** Roadmap of data analysis. **(d)** Dimensionality reduction of the evoked spectral information of all pair-wise evoked potentials in the activated cluster (i.e., pairs with underlying connections). The spectral information is the concatenated power and inter-trial phase coherence (ITPC) spectrograms of the evoked potentials, which are down-sampled to balance the number of datapoints for earlier and later neural responses (see Methods). While this figure is not intended to show distinct clusters, we have colored the dots (a posteriori) to illustrate that data from ipsilateral thalamic stimulations (red dots) are already distinguishable upon a visual inspection. Notably, the color patterns follow different stimulation sites but for recording sites (also see SI **Fig. S3 a,c,d**). They are not biased by specific subjects (SI **Fig. S4**). **(e)** Neural feature encoding involved two main steps: UMAP supervised learning and cluster-based permutation testing. The input data was the evoked power and ITPC spectrograms of the group-level whole-brain evoked potentials, while the dependent variable (i.e., the labels) were the ipsilateral cortical (COR-ipsi), ipsilateral thalamic (THAL-ipsi), contralateral cortical (COR-contr), and contralateral thalamic (THAL-contr) pairs. The algorithm successfully characterized these evoked spectrograms into the four categories. HDBscan was used to formally define the clusters in the embedding space, dissociating them from noisy channels. Original spectrograms of the pairs clustered in the four categories were then used to perform cluster-based permutation testing, by which we identified three time-frequency clusters within the spectrograms that were specific to each category (i.e., Neural Feature). Before statistical testing, the evoked power spectrograms of the four stimulation sites already show distinct features that can be distinguishable by visual inspection alone (marked by white circles and numbered).

As a result, EPs in the activated cluster were differentiated from the spontaneous activity based on prominent changes in both power and ITPC relative to the baseline (pre-stimulation intrinsic neural activities), and also differentiated from the unnatural signals that we have manually labelled as noise (e.g., stimulation artifact or bad channels) by providing those typical cases. Once all subjects’ EPs were successfully classified as “activated” vs. “non-activated”, we ensured the cluster’s meaningful representation of the data by manually inspecting the original and reconstructed data for every subject. With the machine-learning refined labelling (activated vs. non-activated) as a benchmark, the preliminary activation criteria were re-evaluated for further sanity checks (**Table S1**).

We next aimed to characterize the prominent electrophysiological properties of the EPs in the “activated” cluster. Dimensionality reduction with uniform manifold approximation and projection (UMAP) revealed that the inherent structure of the data was different for stimulation-recording pairs within the same hemisphere (i.e., ipsilateral) compared to those across the two hemispheres (i.e., contralateral connections). Surprisingly, we also discovered that the inherent structure of the data was different for EPs generated by stimulation of the thalamic sites vs. cortical sites (**Figure 2d**; more label mapping can be found in **Fig. S3**). The thalamic vs. cortical recording sites, however, do not exhibit such obvious differences (**Fig. S3c** vs. **Fig. S3d**). We refer to this anatomical features (1. Stimulation from the cortex or thalamus, 2. Ipsilateral or contralateral connections) as UMAP localizers. We then searched for the significant features that set the data apart in the embedding space. To dissociate the data that contain the features of interest, we applied supervised learning with the UMAP localizers as labels. As expected, it resulted in four clusters in the embedding space with distinct spectral patterns, i.e., clusters of thalamic, cortical, ipsilateral, and contralateral EPs **(Figure 2e)**.

### Dissociating Distinct Neural Features in Evoked Responses

Once the four clusters were identified, we used the power and ITPC spectrograms from each cluster for further statistical testing. Cluster-based permutation testing identified the significant time/frequency boundaries of three distinct neural features in the data that we labeled as Feature 1 (F1), Feature 2 (F2), and Feature 3 (F3) as tentatively shown in **Figure 2e** by visual identification. Details of statistical testing procedures are presented in the *Methods*. We summarize the three features as follows (**Figure 3a, b**): F1 is characterized by significant increases in power of high frequency activity in the gamma range and the strength of ITPC within the first 10-60ms. Of note, F1 was clearly dissociable from the artifact which was showed as a high-frequency (> 100 Hz) peak power immediately after the stimulation (< 10ms) and lacking signal continuation in both frequency and time dimensions (**Fig. S5)**. F2 is characterizable by an increased power in the theta to alpha range, without phase locking, happening ∼120 ms upon stimulation. Later than F2 (∼200 ms post-stimulation), F3 is featured by a prolonged power increase in the theta band, with a phase-locking feature which indicates oscillation (**Figure 4a**).

**Figure 3.**
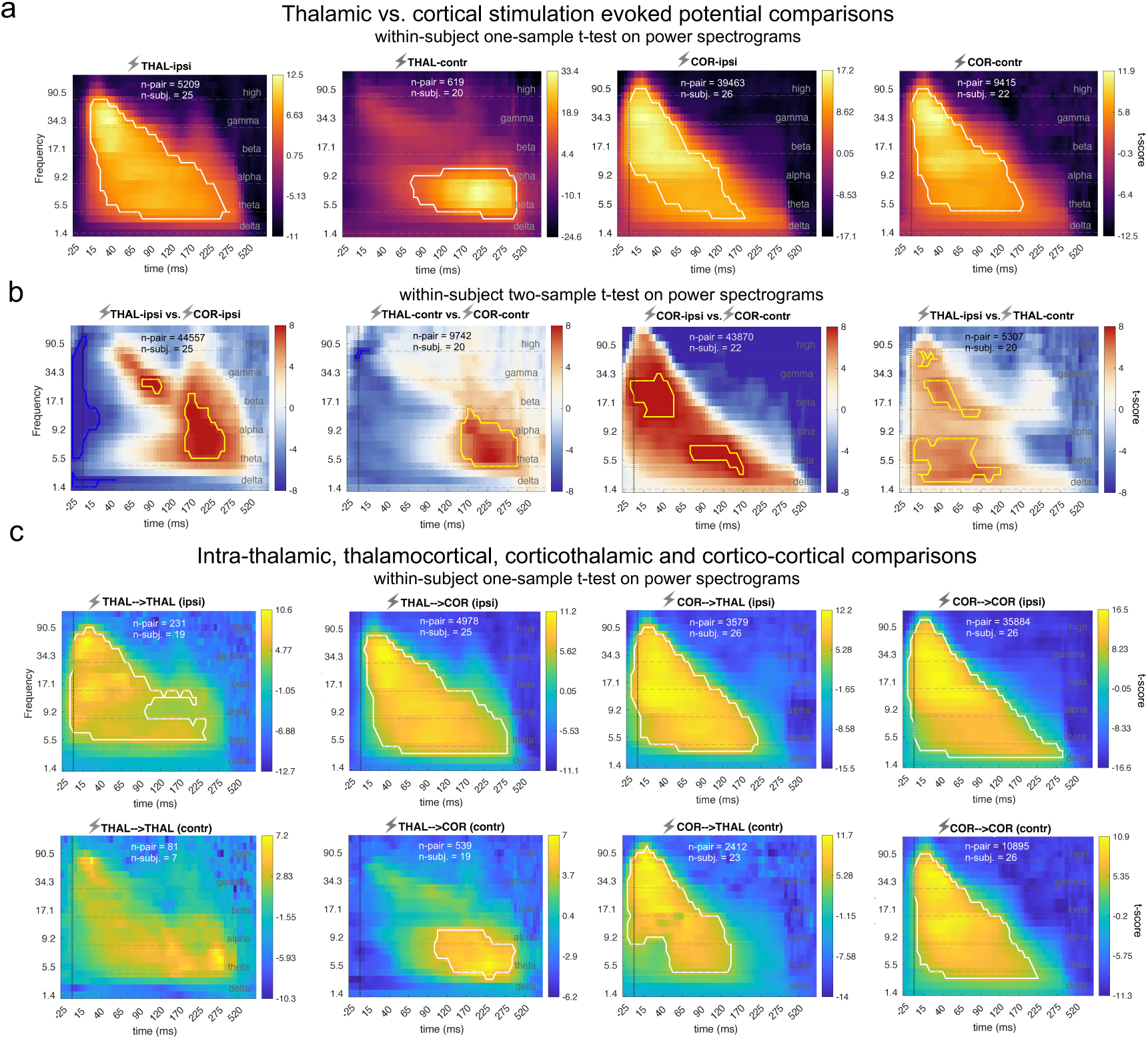
Spectrograms of stimulation-evoked power within and between anatomical categories. (COR: cortex, THAL: thalamus, ipsi: ipsilateral, contr: contralateral, →: causal influence direction). Highlighted contours on the spectrograms indicate significant clusters (n-permutation = 5000, initial cluster forming threshold = 6 t-scores, P_cluster_ < 0.01). Dashed lines on the spectrograms denotes the segments of conventional frequency bands: [0.5, 5] Hz (delta), [5, 8] Hz (theta), [8, 15] Hz (alpha), [15, 30] Hz (beta), [30, 70] Hz (gamma) and > 70 Hz (high). Same tests were conducted on ITPC spectrograms, which generated similar results (**Fig. S6**). The frequency axis is in a logarithmic (log) scale. The time axis is unevenly sampled to balance the varied lengths for different clusters (see Method “training data preparation”). (a) Thalamic and cortical evoked spectrograms and significant clusters in either ipsilateral or contralateral hemispheres. Significance testing conducted using within-subject one-sample t-test (in a mixed-model design): individual-level t-statistics input into a group-level significance testing. The color bars show group-level t-scores. The color bars show group-level t-scores; significant clusters of the one-sided test is marked by white contours. These spectrograms, without significant markers, have also been used in Figure 2e for illustration. (b) Thalamic vs. cortical evoked spectrogram comparison using within-subject two-sample t-test (two-tailed) on the power spectrograms, significance inference performed with cluster-based permutation testing. Yellow and blue contours respectively highlight significant clusters of the contrast indicated on the subtitle and its reversed contrast. Acronym example “THAL-ipsi” on the subtitle means ipsilateral pairs stimulating from thalamus. (c) Intra-thalamic, thalamocortical, corticothalamic and cortico-cortical evoked spectrogram comparisons. Acronym example “THAL-ipsi” means ipsilateral pairs stimulating from the thalamus. Significant clusters of the one-sided tests are marked by white contours. For THAL->THAL (contra) connections, no significant cluster was generated here as there were not enough subjects (n<10) to provide sufficient statistical power for the mixed-model testing. However, a significance test can be done at the level of electrode contacts (i.e., without considering the grouping factor of “subject”). This alternative analysis showed a consistent pattern with the subject-averaged spectrogram, and resulted in two significant clusters, one in the gamma band before 60ms, another in the alpha/theta band around 100-275 ms (**Fig. S8**). Acronym example “THAL-THAL (ipsi)” on the subtitle means ipsilateral pairs stimulating from the thalamus and recording from the thalamus.

**Figure 4.**
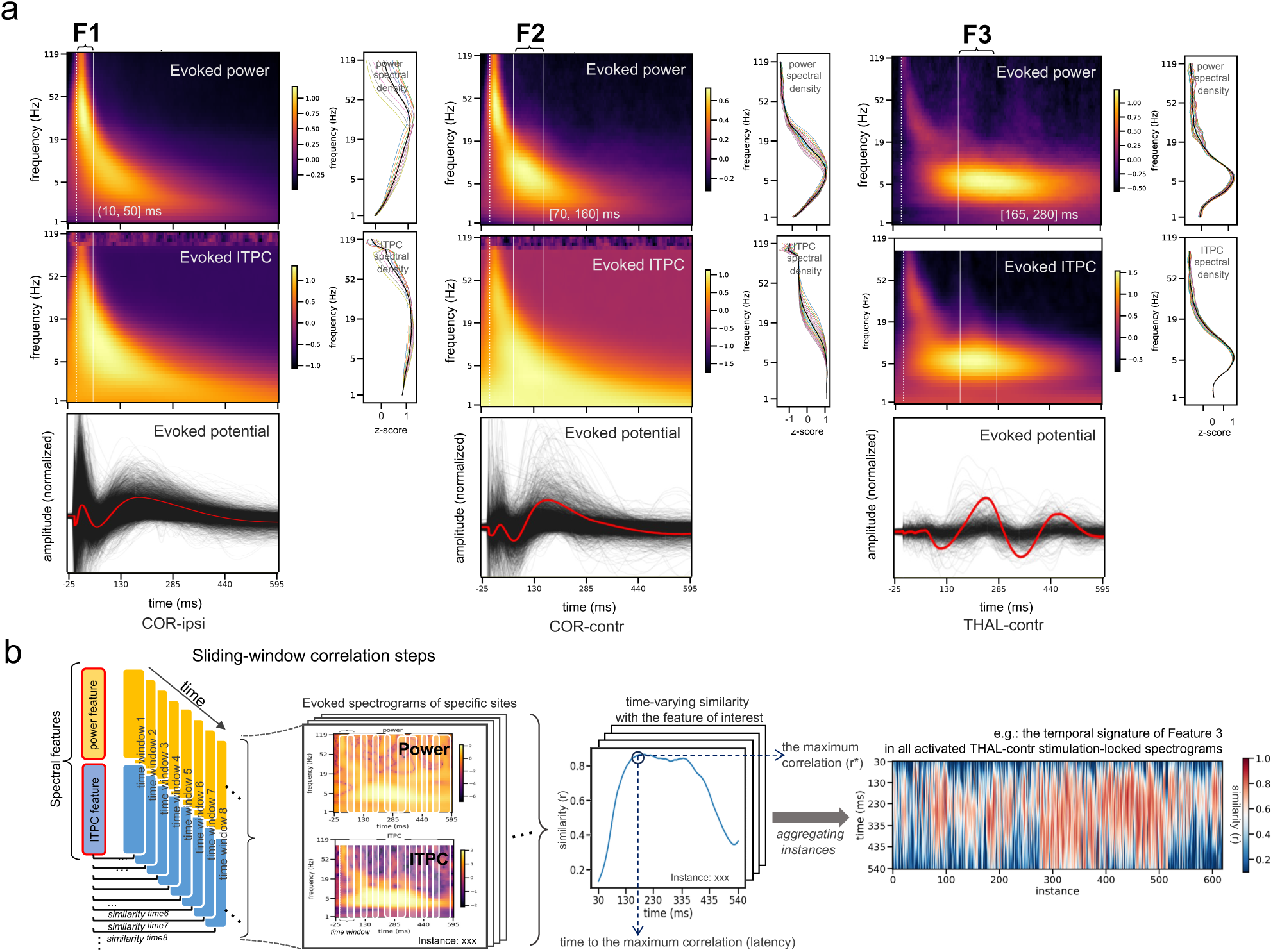
Electrophysiological neural features indicating different types of connectivity and illustration of the decoding process. (a) Neural features specified by the spectra-temporal information. The time windows of the neural features are circumscribed by the significant clusters distinct among conditions from the previous cluster-based permutation testing. The time-frequency relationships in the three features are represented by the group-averaged spectrograms of the COR-ipsi, COR-contr and THAL-contr stimulations, as these three categories show clearest and non-overlapped significant clusters at the group level. The frequency axis is in log-scale. The time axis is in a natural scale with even samples, different from the time axis in Figure 3. The values of power and ITPC spectrograms are transformed (logarithmized and square-rooted, respectively) to approximate Gaussian distributions, and then zscored (in both time and frequency directions) to be comparable among connections. Line plots to the right of the spectrograms show the (normalized and log-scaled) spectral density of the power and ITPC during the significant time window, with colored lines for all the time points, and black for the time average. Since the evoked spectrograms were baseline-corrected, depicted curves can be seen as the “bumps” on top of the 1/f background noise. Group-averaged evoked potentials at the temporal domain corresponding to each category of the spectrograms are shown below. (b) Cross-correlation with a sliding window approach. To reveal the presence of each feature in individual connections, the feature information (i.e. the short-lasting spectral information characterized by the power and ITPC in the time-frequency window) is correlated to the spectrograms with Person-correlation r. To examine this correspondence at every time point (sampling rate is 200 per second), an over-lapping sliding window approach is used, whereby the “transient” time-frequency information at each time point is examined while the examining window is sliding over the spectrograms. This generates the dynamic appearance of the feature presence over time. The line plot in the middle is an example case randomly selected form the THAL-contr instances. Every point on the curve, with a specific time in x-axis and similarity measure in y-axis, indicates how similar the current spectral information (power and ITPC sustaining in certain amount of time) matches the neural feature in concern. The maximum r (r*) and the time to r* (latency) was taken as indices of feature representation and analyzed in the subsequent analyses. The heatmap to the right shows the correlation between all the THAL-contr instances and the Feature 3 (F3). An overview of feature presence in all categories of all features is presented in **Fig. S9**.

Within-subject comparisons showed that the ipsilateral cortical stimulations compared to contralateral cortical stimulations (i.e., COR-ipsi > COR-contr) caused significantly stronger F1 and F2 signals (**Figure 3b**). Similarly, ipsilateral thalamic stimulations compared to contralateral thalamic stimulations (i.e., THAL-ipsi > THAL-contr) caused stronger F1 and F2 signals (**Figure 3b**) i.e., stimulations of the thalamic and cortical sites lead to a stronger F1 and F2 signals in ipsilateral (compared to contralateral) brain sites. Of note, F1 was lacking in the contralateral recordings to the thalamic stimulation (i.e., THAL-contr).

Importantly, the EPs generated from thalamic stimulations were distinct from the cortical stimulations by the presence of a strong F3 signal (i.e., higher theta frequency change starting emerging around 165 ms and persisting for about 250 ms (**Figure 3b**). By checking the signals in the time domain, combining the evidence of the evoked power and ITPC, we confirmed (i) the presence of oscillatory activity embedded in the F3 signal, and (ii) that the F3 signal is not a delayed F2 or continuation of it. In the time domain, F3 indicates a ∼5 Hz oscillation, but F2 indicates a wider curve with jittered peaks across sites. In the time-frequency domain, both F2 and F3 share similar power spectrum, but their ITPC pattern differs significantly in that F2 does not have a phase-locking feature (**Figure 4a**). Moreover, unlike F2, F3 is lacking in cortical EPs, instead, it is specific to thalamic EPs with both THAL-ipsi and THAL-contr clusters showing this delayed and long-lasting F3 activity.

We then investigated the spectrograms separately for different recording sites. Profiles of THAL→COR, COR→THAL and COR→COR in both ipsilateral and contralateral pairs reflected the main results of the UMAP localizers, which showed that the stimulation sites, rather than the recording sites, dominate the spectral features.

Notably, multi-site thalamic coverage enabled us to examine intra- and inter-thalamic causal connectivity at the individual level (**Figure 3c)**. The results provided unprecedent data from the thalamus where we could compare the connectivity of different thalamic subregions with the available brain recording sites and examine the effect of electrical stimulation of a given thalamus upon the activity of sites in other thalamic subregions within the same or across the other hemisphere (**Fig. S7**).

### Timing and Global Patterns of Connectivity Indicated by Feature Presence

Next, we aimed to understand how each of the three neural features unfolds in time within individual instances of connectivity for the four categories of interest (i.e., COR-ipsi, COR-contr, THAL-ipsi and THAL-contr). The group-averaged spectral information within the significant time windows from the COR-ipsi, COR-contr and THAL-contr categories was used to create the feature template for F1, F2 and F3, respectively (**Figure 4a**). Using a sliding-window correlation approach (see details in *Methods and* illustration in **Figure 4b**), we estimated the strength of feature representations (r*) and their presence in time.

Corresponding to the previous group-level cluster-based permutation testing on the spectrograms, the decoding results also showed that both F1 and F2 peaks were seen more strongly for ipsilateral than contralateral EPs (Δr* = 0.11, t = 48.05, p_FDR-corr_ << 0.01, n-connection [conn.] = 57881, n-subject [sbj.] = 26 for F1, and Δr* = 0.09, t = 44.90, p_FDR-corr_ << 0.01, n-conn. = 57881, n-sbj. = 26 for F2, using the stimulation category “COR/THAL” as a covariate, corrected for the nested individual, regional and site effects in a hierarchical linear model [HLM]). Additionally, the decoding method was able to provide more precise time information than the previous test, which showed that both F1 and F2 were detected earlier in the EPs responding to cortical versus thalamic stimulation (Δlatency = 5.60 ms, t = 27.20, p_FDR-corr_ << 0.01, n-conn. = 37444, n-sbj. = 26 for robust F1 [r*>0.4, latency in the 10-100 ms range], and Δlatency = 8.18 ms, t = 22.49, p_FDR-corr_ << 0.01, n-conn. = 48300, n-sbj. = 26 for robust F2 [r*>0.4, latency in the 70-200 ms range]; using the cross-hemispheric category “ipsi/contr” as a covariate in an HLM).

The decoding results also echoed the previous finding that F3 was seen mainly in evoked potentials pertaining to the thalamic stimulations, with F3 peaking higher in ipsilateral than in contralateral EPs (Δr* = 0.11, t = 48.05, p_FDR-corr_ = 0.02, n-conn. = 6497, n-sbj. = 25), but without significant difference in the latency between ipsilateral and contralateral EPs (Δr* = 5.03 ms, t = 1.86, p = 0.06, n-conn. = 3106, n-sbj. = 25 for robust F3 with r* > 0.5, latency > 200 ms). Additional post-hoc comparisons among sub-groups are presented in **Figure 5** and SI **Table S9**. The correspondence of the decoding results with the previous group-level spectrogram analysis suggests that our devised feature indices can reliably indicate the presence of the multivariate features with single values.

**Figure 5.**
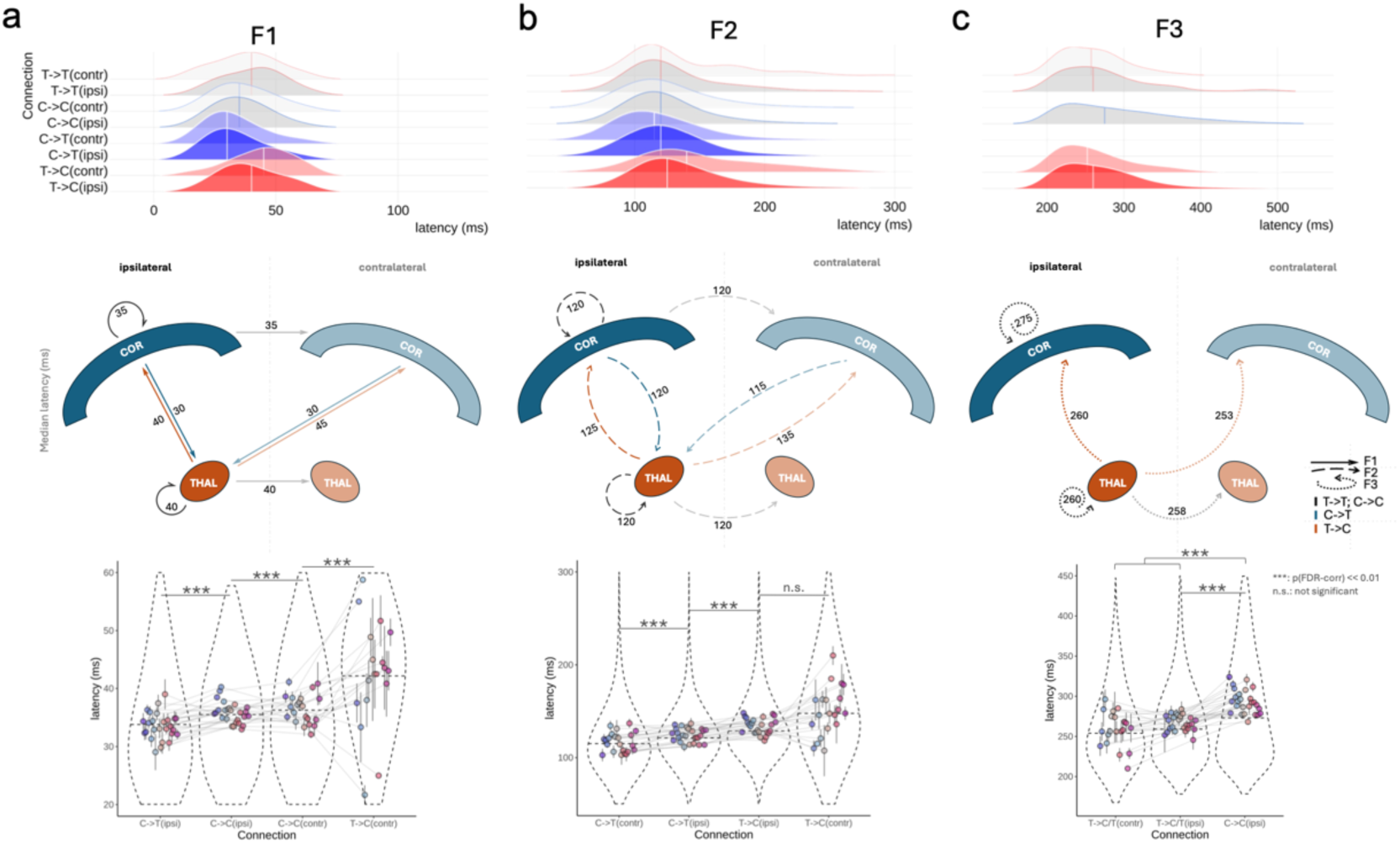
The timing of feature presence (i.e., latency of maximum representation of the feature). (a) The top panel shows the distribution of the latencies of F1 presence in all types of connections. F1 is identified in individual instances by finding the peaks of the feature curve. Included in the distribution are connections with strong F1 representation (r* > 0.4) which falls in a sensible latency range of (10, 70) ms, based on our results. Density distributions for thalamocortical connections are filled red, corticothalamic connections are filled blue, and thalamo-thalamic/cortico-cortical connections are filled grey. Colors for contralateral connections are lighter than those for ipsilateral connections. The middle panel is a cartoon showing the median latency of the feature presence among the aforementioned connections. The lower panel depicts within-subject post-hoc comparisons (adjusted for individual and regional differences) between the categories that showed evident differences in the latency distributions. Each colored dot marks the mean latency of all the connections from one subject. The black bar through the dot marks the range of ± one standard error of the within-subject data. Grey lines across groups show the within-subject comparisons. All tests were corrected for multiple comparisons. The full list of post-hoc comparisons is presented in SI Table S9. (b) and (c) shows F2 and F3 results with same visualization schemes. The criteria for data going into the F2 distributions are r* > 0.4 and latency in (70, 165) ms. The criteria for the F3 distributions are r* > 0.5 and latency in (200, 400) ms. As there were much fewer contralateral thalamocortical connections with F1 and F2, the data for these two features showed large variabilities.

We further examined the whole-brain spatiotemporal patterns indicated by the three neural features. To do so, we constructed connectivity matrix with each cell filled with the connectivity index of R for indicating the signal strength between two brain regions (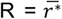, averaged across the sites where the feature is present in the given regions). By organizing brain areas in the order of anatomical proximity, interesting patterns emerged: (i) the F1 matrix demonstrated a profile of modular connectivity, i.e., higher F1 representation was seen for connections between pairs located within the same hemisphere or lobe and within the ipsilateral thalamus (**Figure 6a, b**); (ii) the F2 matrix revealed wide-spread representations across regions and hemispheres and demonstrated that the global architecture of connections based on F2 did not differ much from the F1 matrix. This seems to suggest that F2 is a combination of jittered rebounds following the same paths for the F1 signals (**Fig. S10**); (iii) unlike the F1 and F2 matrices which have a modular architecture, the F3 was sparsely present among the cortical connections, but largely present in the cortical responses (i.e., widespread and bilateral) associated with thalamic stimulation (**Figure 6f, g**).

**Figure 6.**
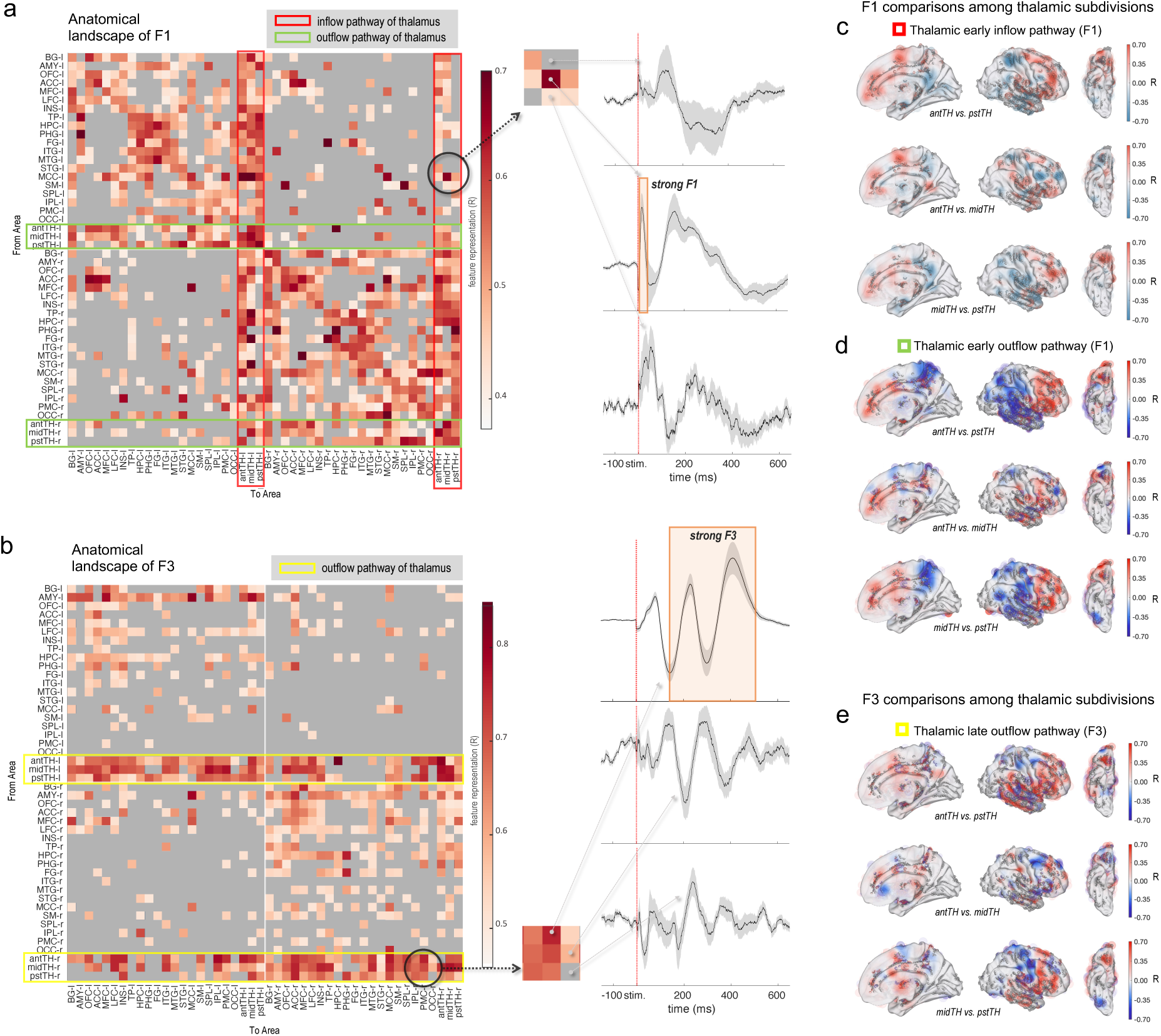
The anatomical landscapes of feature presence. (a) Early F1 connectivity matrices for bilateral cortical and thalamic subregions (l: left, r: right, ant: anterior, pst: posterior, PMC: posteromedial cortex, SM: sensorimotor, SPL: superior parietal lobule, ACC: anterior cingulate cortex, CLT: claustrum, TH: thalamus, IPL: inferior parietal lobule, MFC: medial frontal cortex, OFC: orbital frontal cortex, STG: superior temporal gyrus, LFC: lateral frontal cortex, INS: insula, FG: fusiform gyrus, HPC: hippocampus, MTG: middle temporal gyrus, PHG: parahippocampal gyrus, AMY: amygdala, TP: temporal pole, MCC: midcingulate cortex, ITG: inferior temporal gyrus); The anatomical localization of all electrode contacts was visually inspected and labeled by an experienced neuroanatomist using the position of the electrodes in the individual subject’s native brain space. Each entry of the matrix has a row and column identity corresponding to site of stimulation and recording, respectively, and its value indicates the feature representation (R) for the corresponding feature. Specifically, 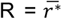, was averaged across connections where the feature was present. Feature presence was binarized with a threshold of r*>0.4 for F1,2 and r*>0.5 for F3. Arbitrary thresholding was applied only for visualization purposes; un-thresholded matrices are presented in the SI **Fig. S10**. From the matrix, three entries were randomly chosen, with graded R values from low to high, to demonstrate their associated evoked responses in the time domain. For plotting the evoked responses, the black line is the site-averaged signal across the evoked potentials stimulated/recorded in the same anatomical regions – it is a group-level average which may involve different subjects whose stimulated/recording sites are in the same brain region. The grey-shaded area around the line is the standard error over sites. (b) Delayed F3 connectivity matrices for bilateral cortical and thalamic subregions with the same visualization scheme as aforementioned. (c) and (e) show brain heatmaps of the contrasted feature representation among the thalamic divisions (antTH vs. pstTH, midth vs. pstTH, midTH vs. pstTH), respectively for the thalamic inflow (i.e., recording in the thalamus) and outflow (i.e., stimulation in the thalamus) pathways. All values were projected to one hemisphere for compact visualization. FS_LR brain surface space (with symmetric left and right hemisphere) is used to minimize the visualizing bias caused by interhemispheric anatomical differences (https://osf.io/k89fh/wiki/Surface/). Grey dots on the brain surface indicate electrode coverage, and the color radius around the dots indicate local R values of the given contact. Due to sparse recording, the color is also projected to the brain surface with a Gaussian function over distance from the source, to approximate a whole-brain level estimation. Coloring intensity on the brain surface has been adjusted for regional density of electrode coverage. The presented brain heatmaps are not thresholded; formal statistical testing results and model details can be found in **Table S2-5**.

In a closer examination, we found that F1 corticothalamic and F3 thalamocortical connectivity features had the following unique characteristics in common: (i) F1 responses recorded in the thalamus had widespread and bilateral origins across the brain (i.e., stimulation of many cortical regions bilaterally could evoke responses in the thalamus) (**Fig. 6a**), and vice versa for F3, the stimulation of a thalamic site would cause widespread cortical areas bilaterally to show delayed slow oscillations (**Figure 6b**); and (ii) both corticothalamic F1 and thalamocortical F3 responses were *stronger* and *earlier* than corticocortical F1 and corticocortical F3 responses, respectively. The statistical details are presented as below.

We compared the cortical versus thalamic targets responding to the same cortical seed of stimulation in the same participant, and confirmed the F1 responses in the thalamus were *earlier* and *stronger* (Δlatency = 2.79 ms, t = 12.74, p_FDR-corr_ << 0.01, n-conn. = 48446, n-sbj. = 26; Δr* = 0.06, t = 18.20, p_FDR-corr_ << 0.01, n-conn. = 30965, n-sbj. = 26, with HLM corrected for the covariance effect of cross-hemispheric connections and the nested random effect of 1378 different stimulation sites of different brain areas in all subjects) (**Figure 6a** for R matrix, SI **Fig. S10** for latency matrix). In other words, stimulation of a given cortical site changed the activity of the thalamus before it changed the activity of its cortical targets (**Figure 5a** and SI **Table S9** for post-hoc latency comparisons).

Moreover, the F3 matrix revealed that the stimulation of thalamic sites caused delayed low-frequency oscillatory effects (i.e., F3) in a large mantle of the cortex in not only ipsilateral but also contralateral hemispheres (**Figure 6b**). By contrast, the F3 feature was only sparsely present with the stimulation of non-thalamic seeds. Importantly, the latency of F3 responses caused by thalamic stimulations was significantly earlier than the non-thalamic seeds (36.27 ms earlier, t = 21.60, p_FDR-corr_ << 0.01, n-conn. = 20823, corrected for the covariance of cross-hemispheric effect, and the nested random effect of 3041 recording sites of different areas in all subjects with an HLM) (**Figure 5c**).

### Differential connectivity among thalamic subregions

Up to this point, we have remained agnostic to the specific anatomical location of the thalamic sites where evoked responses were recorded or stimulations were seeded. However, each subject had more than one electrode in the thalamus targeting the anterior, mediodorsal and pulvinar nuceli^5^. Given that the precise anatomical boundaries of each thalamic nucleus within each subject may be variable and difficult to ascertain with neuroimaging data, and that the field of evoked responses or electrical stimulations may cross the nuclear boundaries, we refrained from labeling our thalamic sites according to specific nuclei, and instead, referred to them as anterior (antTH), middle (midTH), and posterior thalamus (pstTH).

Leveraging the anatomical information of recordings and simulation sites - at the individual level - we compared the connectivity profiles of the three different thalamic sites. This analysis revealed unique site-specific profiles of connectivity between the thalamus and the cerebral cortex (**Figure 6c, d, e)**. As describing the details of differential connectivity of thalamic sites are outside the scope of this report, we only highlight some of the most salient findings. The full results of pair-wise comparisons among the thalamic subregions for their bi-directional connections with each of the brain regions are provided in the supplementary **Table S2-7**.

Next, we compared the delayed thalamocortical F3 signal representation map with the ones for early thalamocortical F1 and early corticothalamic F1 maps, with pair-wise comparisons within the same pair of thalamic vs. non-thalamic sites. Using the measure of normalized feature representation 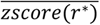 as shown in **Figure 7** and detailed in **Tables S6 and S7**, between a subregion of the thalamus and a given brain region of interest, we found significant asymmetry between thalamocortical F3 representations and thalamocortical F1 representations. Many regions had stronger F3 than F1 representations suggesting that the delayed thalamic (F3) outflow is represented across a more widespread cortical areas compared to the fast thalamic (F1) outflow signal (see brain regions with more brown colors in **Figure 7a** and positive estimate numbers in **Table S6**). The same applies to the comparison between thalamocortical F3 and corticothalamic F1 representations. In this comparison, the hippocampus (HPC) was an exception among all other examined brain regions: representation of F1 signals evoked by HPC and recorded in anterior and posterior thalamic sites was stronger than the representation of F3 signals recorded in HPC and evoked by the thalamus, with pair-wise comparisons for the same sites in the regions of interest (Δz = 0.30, t = 2.51, p_FDR-corr_ =0.04, n-conn. = 190, n-sbj. = 21 for antTH; Δz = 0.62, t = 4.72, p_FDR-corr_ << 0.01, n-conn. = 194, n-sbj. = 18 for pstTH) (**Table S7**).

**Figure 7.**
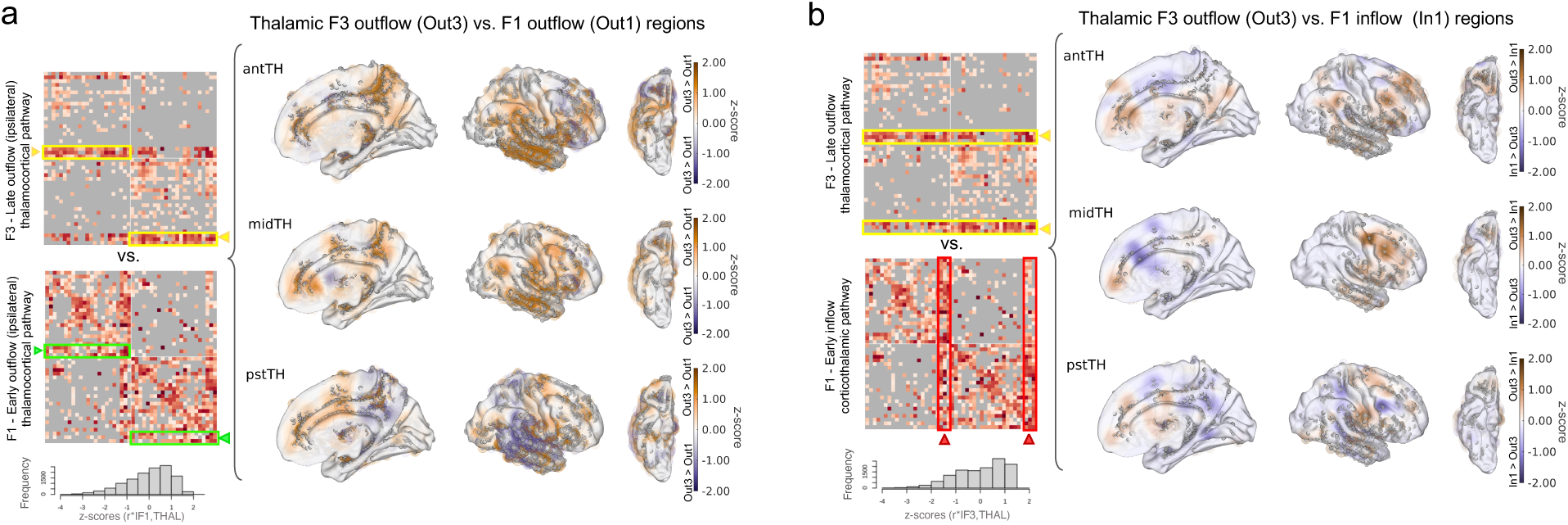
Late thalamocortical compared with early corticothalamic connections. Causal connectivity matrices are adapted from Figure 4 where the details can be found. These are shown again to indicate the data used for the respective brain heatmap plots. (a) The brain heatmap shows the comparison between F1-outflow vs. F3-outflow representations measured in each recording site across the brain due to stimulation of the anterior, mid-, and posterior thalamic subregions corresponding to anterior, mediodorsal, and pulvinar nuclei of the thalamus. Since we did not have hypotheses about hemispheric lateralization of connectivity profiles, we projected all electrodes onto one hemisphere for compact visualization. FS_LR brain surface space (with symmetric left and right hemispheres) was used to minimize the visualization bias caused by interhemispheric anatomical differences (https://osf.io/k89fh/wiki/Surface/). The color bar shows normalized feature representation: 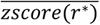, averaged across the stimulation sites Before averaging, the r* values of all the thalamic connectivity with the same feature type were z-scored, i.e., *zscore(r*|F1,THAL_inflow&outflow_), zscore(r*|F3,THAL_inflow&outflow_)*, in order to make the feature representations between pathways/feature types comparable. The distributions of these normalized *z*-scores are presented below the connectivity matrices. The positive (brown) patches represent contrasted z-scores of F3-outflow (out3) being greater than F1-outflow (out1), while blue patches represent brain regions where F1 outflow (out1) is greater than F3 outflow (out3) representation. To avoid negative values being subtracted to become a positive value, negative z-scores were zeroed before being contrasted. Grey dots on the brain surface indicate electrode coverage. (b) Same visualization scheme for the comparison of F3-outflow vs. F1-inflow pathways of the thalamus. The presented brain heatmaps are not thresholded; formal statistical testing results are presented in supplementary **Table S6-7**.

## Discussion

In our study, we addressed the limitations of past studies by leveraging a novel approach to characterize the complex features of electrophysiological responses that are evoked by not only cortical but also thalamic stimulations in the human brain. This is important because, as we documented in Fig 1, reliance on the magnitude of recorded neural responses or expecting typical N1 and N2 profiles in the evoked responses may greatly bias our views of causal connections in the brain. The magnitude of evoked responses can be affected by several confounding factors such as distance from the source of stimulation, volume conduction, and electrode impedance^12^ and the time-to-peak may be elusive when there is no clear N1 or N2 peaks. While typical N1 and N2 waveforms reported in the CCEP literature are abundant, it is important to note that the past reports were based on group level analysis of data from cortical stimulations^13,14^. As noted in **Figure 1**, individual evoked signals may be far more complex than simple N1 and N2 waveforms and the magnitude of evoked responses may not always pass an arbitrary threshold we set for significance. This issue is even more pressing when waveforms from subcortical structures such as the thalamus are included. Therefore, in line with recent attempts by other investigators^7,8^, we felt it was imperative to employ a multivariate and unbiased approach for sorting the distinct profiles of connectivity across cortical and thalamic sites.

In our encoding method, we relied on the machine learning algorithms of UMAP to identify the inherent variabilities among the datasets. We emphasize that our data-driven approach was not blind to biologically plausible features. Instead, it was directly guided by the 2/3 of the data that had already been labelled based on established neurophysiological criteria for significant evoked responses (i.e., significant amplitude above the noise level, significant inter-trial phase coherence concerning reliability of responses across trials, and more importantly, labeling what constitutes a stimulation artifact). Our post-hoc analysis also confirmed that the machine-learning refined labeling largely corresponded to the activation detection results based on the preset biological criteria, among which, the criterion based on significant ITPC (i.e., consistency in the evoked responses over trials) had the highest correspondence (0.80). In contrast, the traditional activation detection criterion of large peak detection (> 5 z-score) had a high precision (0.99) but low sensitivity (0.53), which is in line with our observation: many thalamic evoked responses (especially the contralaterally recorded ones) would be missed if based solely on the amplitude criterion as those responses usually have lower amplitudes on the voltage traces, though not necessarily possessing less robust evoked features.

Our new approach yielded important results suggesting that the complex electrophysiological signals evoked by repeated single pulse electrical stimulations have three distinct features (i.e., F1, F2, and F3). In the context of past literature, unitary F1 and F2 features may correspond to clear N1 and N2 peaks, but we emphasize that changes in the neural activity of a given brain site evoked by the stimulation of another site cannot always be depicted as a uniform waveform consisting of N1 and N2, but rather, a complex signal with different components of the evoked waveform which may in turn reflect distinct physiological substrates with distinct sources. While we postulate that F1 and F2 may be related to the well-known N1 and N2 components of the EP signal described in the literature ^12,14,15^, our data suggests that F3 is a unique feature of its own and not a prolonged F2 – even though F2 and F3 features have seemingly similar power spectrogram peaking at a low frequency range. Our rationale is as follows: (i) F3 has a peak oscillation at about 5 Hz, but F2 has wider frequency range with jittered peaks across trials. (ii) F3’s ITPC spectrogram differs significantly from the F2’s, as F3 has a phase-locking feature at the theta frequency but F2 does not (**Figure 4a**). The oscillatory feature of F3 was also observed at the same time-frequency cluster as long-lasting theta oscillations (>= 2 cycles). The oscillatory feature of F3, unlike F2, was observed in the time domain: it was strongly time-locked and exhibited at least two cycles of oscillation - even when averaging the EPs recorded across all recording areas and across all thalamic stimulation sites. (iii) The temporal window of significant time-frequency clusters in F2 and F3 do not overlap (i.e., F3 always happens after F2, peaking > 200 ms post-stimulation). (iv) Lastly, F3 feature is present specifically when the thalamus is stimulated (including both THAL-ipsi and THAL-contr clusters), while F2 does not have this anatomical specificity.

We documented that the F3 feature was detected with the stimulation of *all three thalamic sites*. This suggests that the F3 signal is not produced by the stimulation of only one specific thalamic nucleus. However, it’s important to acknowledge a limitation of our study, which stemmed from the absence of data from a number of other thalamic nuclei. For instance, we have no information about the profile of electrophysiological responses evoked by the stimulation of thalamic nuclei such as the geniculate bodies. This limitation was primarily due to the clinical constraints and ethical considerations inherent in studies involving human subjects.

We also note that the F3 oscillatory evoked responses were also seen with the stimulation of a few non-thalamic brain regions - especially the hippocampus and amygdala – both of which are non-cortical structures [**Figure 6b**]). interestingly, the latency of F3 signal evoked by the stimulation of these non-thalamic sites was significantly delayed compared to the latency of F3 signal evoked by the stimulation of the thalamic sites, and they were mostly present in ipsilateral recording sites while the thalamic stimulations evoked F3 in far more widespread areas involving bilateral hemispheres.

As the thalamus is considered to play a key role in cortical rhythms^16^, and as the hippocampus is known for its role in generating theta rhythms^17^, we hope our findings will motivate future systematic studies to determine how theta oscillations observed throughout a broad surface of higher association areas in the cerebral cortex during cognitive activities tasks or rest^18,19^ are related to the delayed modulatory effects exerted by the thalamic (or medial temporal lobe) structures. It also remains to be determined if the bilateral presence of the F3 signal evoked by thalamic stimulations is functionally related to, or is crucial for, the integration of information across the two hemispheres.

Our approach with signal decoding of the stimulation and recording data revealed that stimulation of a given cortical region evoked *earlier* and *stronger* responses in the thalamus compared to the cortical targets. A hypothetical interpretation of this finding is that the thalamus acts as a universal receiver of information being exchanged across cortical regions (i.e., a “listener of corticocortical dialogue”) – which is a necessary for it to function as a key structure for information integration as detailed recently^2^.

Additionally, the multisite thalamic stimulation and recording within the same individual offered us an unprecedent opportunity to examine the intrathalamic and interthalamic causal connections. We demonstrate that the stimulation of a given thalamic site led to fast high frequency modulation of activity in other thalamic sites ipsilaterally as well as delayed low frequency modulation of thalamic sites contralaterally. These findings provide a plausible electrophysiological mechanism for intra-thalamic and inter-thalamic connectivity, which may have important implications and clinical relevance: Stimulating a specific thalamic nucleus with DBS (e.g., for treatment of epilepsy^20^) is likely to impact the activity of other thalamic nuclei (beyond the one directly targeted) and the cortical networks they are associated with. Thus, the therapeutic benefits of DBS might be related to a broader network of brain regions that are modulated rather than a focal network targeted. However, this does not imply that the effect of stimulation of each targeted thalamic site is diffuse and all-encompassing, as shown in **Figure 5**, stimulations in three subregions of the thalamus do not elicit anatomically identical responses, but instead show relative anatomical preference.

We are mindful that our data was acquired from patients with a neurological disease (epilepsy) and one may ask to what extent they are generalizable to normal human brains. While we acknowledge that molecular and cellular changes accompany local circuit rewiring in patients with chronic epilepsy^21^, there is no firm evidence suggesting changes in the fundamental topology of connections at the system level in these patients^22^. In the extant literature, unless studying the circuit level and local changes in a region-of-interest type of analysis^23–25^ or graph theoretic measures for deriving high-level estimation of network capacity^26,27^, the global network topology of the brain has been shown not to be significantly different between epileptic cohort and the normal control^28^. Several studies focusing on epileptic brain reorganization have shown that the same sets of canonical brain networks can be discovered by the data-driven methods, just as in normal controls^29–31^.

We are also mindful that electrode coverage in our subjects was limited and sparse. This is unfortunately a limitation of the intracranial approach in human participants that we, as researchers, have little control over since the implantation of electrodes in all participants was clinically driven. We made the best effort to reduce the confounding effects by maximizing the number of subjects, and diversifying the cohort with different epilepsies so as to avoid a system bias of encountering pathological tissues concentrated in specific areas.

Despite the limitations of our study, we hope that our findings can serve as the steppingstone towards a wider understanding of the architectural topology of interactions across the cerebral cortex and between those and subcortical structures such as the thalamus.

In closing, we acknowledge that the causal connectivity maps presented in our current study provide only one aspect of the functional architecture of the brain, and as such, they serve as rough approximations for the dynamic causal interactions occurring in real-life scenarios and during cognitive processing and are complimentary to those connectivity measures generated by structural diffusion MRIs or resting state or task specific imaging studies^3,4,32–34^. To highlight the functional significance of RSEPS-based connectivity maps, we recently confirmed that the strength of RSEPS connectivity between sites correlates with the strength of their co-activations during a cognitive task, i.e., if a specific pairs of neuronal populations across the cortex and thalamus had high RSEP-based connectivity strength they had higher correlation of high-gamma activations during a memory task^11^. Additionally, it has been shown that seizures propagate from cortical seizure onset zones to thalamic sites^5^ or other cortical areas^35^ that show faster and stronger RSEPS connectivity measures with the seizure onset zone.

## ONLINE METHODS

### Resource Availability

#### Materials availability

Raw data, including electrophysiology collected in this study will be shared publicly after publication.

#### Data and code availability

- The electrophysiological data have been deposited at Mendeley and are publicly available as of the date of publication. The DOI will be listed in the key resources table.
- All original codes have been deposited at Zenodo and is publicly available as of the date of publication. DOIs will be listed in the key resources table.
- Any additional information required to reanalyze the data reported in this paper is available from the lead contact upon request.

### Participants

In this study, twenty-seven participants (40.7% female; mean age ± SD: 34.9 ± 10.0 years) diagnosed with focal epilepsy were recruited (see **Table 1**). Each participant had 180 ± 46 (mean ± standard deviation) SEEG contacts implanted. The total number of electrode contacts being studied is 4864. All participants underwent invasive electrophysiological monitoring at our medical center as part of their treatment for refractory epilepsy. The placement of electrodes was determined exclusively based on clinical requirements. Prior to their involvement, all participants provided written informed consent, and the study protocol was approved by the Institutional Review Board for human experimentation.

**Table 1.**
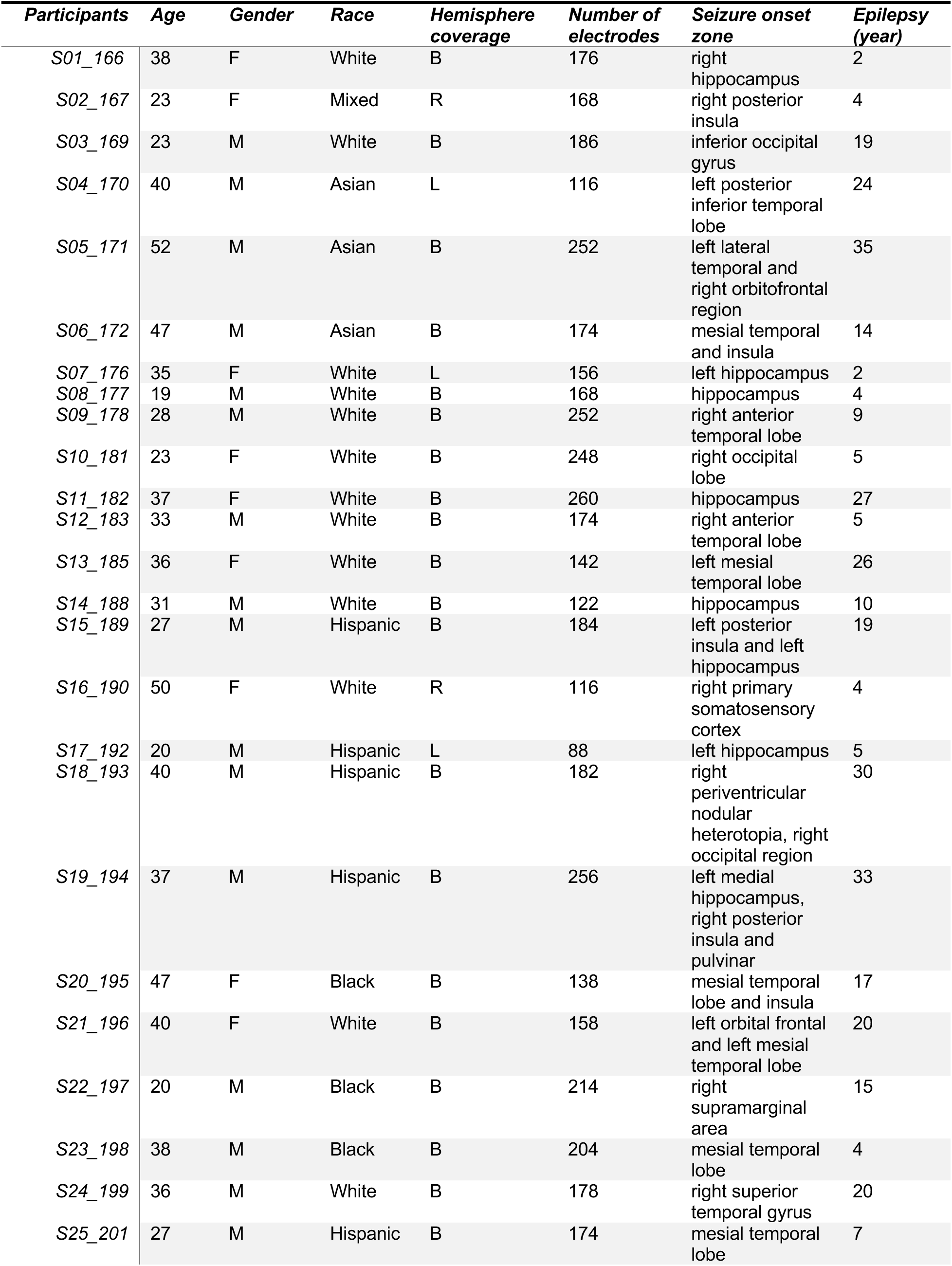
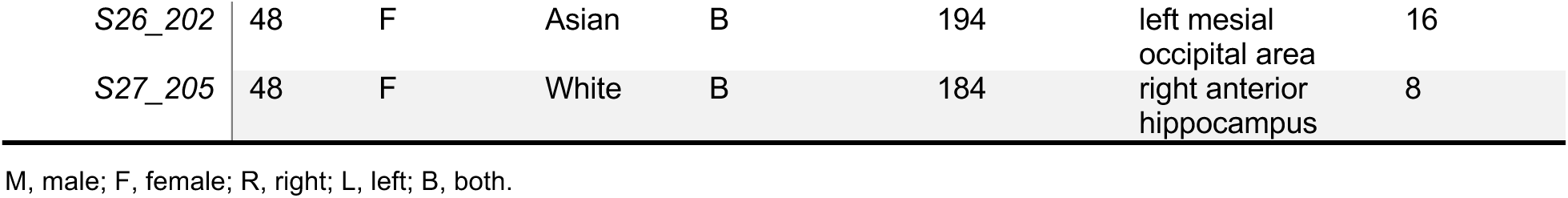
Participant demographics.

### Patient Safety & Ethics

As noted, we did not implant extra electrodes for thalamic recordings. We only extended the electrodes that were clinically planned for cortical monitoring to reach the thalamus. Our procedure was based on patients’ informed consent and all our research related activities were IRB approved. To minimize injury, we used a reduced diameter obturating stylet and reduced diameter electrodes with 0.86mm diameter (AdTech, Inc). With this surgical approach, we have observed no complications, no thalamic hemorrhage or edema, and no neurological symptoms post operatively in any of the patients examined. We note that the thalamic recording in epilepsy patients has been widely practiced in some centers in Europe for two decades yielding clinically important information^36–40^. The method has recently gained ground in the U.S.^41–44^, and a recent neurosurgery editorial recommended and encouraged more centers in the U.S. to do the same^45^. Additionally, iEEG implantation strategy is decided by a group of clinicians who meet weekly to decide the clinical needs of the patient, and not by this research project. Also, as noted, all patients provide informed consent prior to being enrolled in any research studies.

### Electrode Implantation

MRI sequences for imaging the thalamus and its subnuclei included BRAVO, fGATIR and MP2RAGE. We co-registered the post-implant CT with a pre-implant MRI. High resolution T1, fast gray matter acquisition T1 inversion recovery (FGATIR), and T1 post-contrast imaging were used for planning. Trajectories were planned to traverse in an orthogonal plane to capture the cortical frontal or temporal operculum, insula, and to be extended into specific sites of interest in the thalamus. The priority for all trajectories was safety and avoidance of blood vessels. To achieve implementation of our multisite sampling approach, the cortical electrode trajectories were extended to reach three different thalamic sites: anterior, mid, and posterior thalamus as detailed in our recent publication^5^. Approximate locations and number of electrodes along with their trajectories were all planned in a multidisciplinary surgical epilepsy conference with detailed review of presurgical data leading to a clinical hypothesis of the most likely seizure onset zones. Electrode placements were at the discretion of the clinical care team, who were not specifically involved with this research project.

### Co-localization of electrodes

A thin cut head-CT, obtained after electrode implantation, was co-registered to pre-operative 3Tesla MRI data for verification of the trajectory. Precise electrode positioning in the FreeSurfer surface space, voxel space and MNI space were automatically extracted by the iElVis toolbox^46^. The T1-weighted MRI scan was used to generate 3D cortical volume and subcortical segmentation using recon-all command of Freesurfer v7.3.2^47^. The post-implant CT scan were co-registered to the pre-implant MRI using the *flirt* from the Oxford Centre for Functional MRI of the Brain Software Library^48,49^ or using *bbregister* from Freesurfer^50^ to get the best results. We manually labelled each electrode on the T1-registered CT image using BioImageSuite^51^. The electrode coordinates in the native anatomical space were inspected and manually labeled by the experienced neurologist J.P., based on the individual brain’s morphology and landmarks.

### Thalamic parcellation

To check the concordance between manual and automatic labeling of thalamic electrode sites, individual contact centers of electrode masses were defined in native T1 space for each subject. The center of mass of each contact was then converted into an X,Y,Z coordinate in the MNI space. A recently developed and widely used MNI thalamic atlas^52^ was utilized to parcellate the nuclear location of each contact within the thalamus. For this, a 1mm cubic voxel region was created around each contact center of mass (contact neighborhood). For each contact neighborhood, the fraction of voxels that overlapped with each thalamic nucleus in the THOMAS atlas was calculated. Contact neighborhoods that fell solely within a single THOMAS nuclear mask would have a value of 1 for that specific nucleus, while contact neighborhoods that had no overlapping voxels with a given nuclear mask would have a value assigned to 0 for that specific nucleus. Thus, for every contact neighborhood anatomically inside the thalamus, the fractional overlap with each THOMAS nuclear mask was calculable. For each site, this within-subject fractional overlap was summed across subjects and across all thalamic contacts to generate overall contact neighborhood fractional overlap values.

### Identifying Regions of Interest

All electrodes contact locations were manually labelled with brain regions of interest (**Figure 1a**) by the experienced neurologist J.P. on the team. This process involves carefully inspecting the CT-MRI fused images in axial, sagittal, and coronal planes for the implanted electrodes, and their surrounding anatomical landmarks in the brain morphology (e.g., basal ganglia including putamen, and pallidum etc.). Contacts entirely or partially located in the white matter (e.g., internal capsule, or cingulum etc.) were labelled as white matter. We emphasize that these regions of interest were not identified in the standard atlas space, but in each participant’s own high-resolution T1 scan to ensure anatomical precision of electrode localization.

### Repeated single electrical pulse stimulation (RSEPS)

Stimulations were performed in a bi-polar manner stimulating between a pair of electrodes. Each RSEPS trial entailed an instantaneous (pulse duration = 0.2 ms) 6 mA biphasic square-wave pulse. For electrode contacts near the seizure onset zones (identified by the clinical team), we stimulated at 4 mA. Repeated pulses (n = 42 ± 2 trials) were delivered in a row with an interval of 2 seconds. We excluded data from (i) bipolar channels with both contacts located in the white matter (see above for details of anatomical localization), (ii) bipolar contacts not within the same anatomical region (e.g., one contact in insula and the other in orbitofrontal cortex), (iii) recording bipolar sites too close to the stimulation site (Euclidean distance between their middle points <= 5 mm), and (iv) channels largely contaminated by stimulation artifact, i.e., recording large signals (>= 7 SD of the baseline) only in the artifact window (< 10 ms), and (v) bad channels based on more specific criterion as detailed below.

### Intracranial EEG data acquisition and preprocessing

The data was collected using the Nihon Kohden recording system with a sampling rate of 1000 Hz. We used an in-house programmed preprocessing pipeline to denoise the intracranial EEG data before any statistical testing. The pipeline included notch filtering at 60, 120 and 180 Hz, data exclusion, and re-referencing. Excluded data includes pathological, noisy channels, and bad trials, the identification criteria of which are described as follows. We used a recently developed algorithm to identify pathological channels by the presence of pathological high-frequency oscillations (HFOs) ^53^. Noisy channels were identified by the presence of extremely high raw amplitude (> 5 standard deviations [SD] across all channels) and/or the prevalence of spikes (> 3 median of the distribution across all channels), i.e. jumps between consecutive data points larger than 80 μV. Trials with extreme variations in the signal, i.e. having extreme mean (> 4 SD across all trials) of the absolute magnitude of the signal values, and extreme standard deviation (> 3.5 SD) of the variance of the signal values along the sampled timepoints, were excluded for being trial-averaged to create the time-domain evoked potential. Following data screening, we applied bi-polar re-referencing instead of common averaging, given that the stimulations were performed in a bi-polar manner, i.e., injected between a pair of electrodes. Common average reference of all non-noisy channels for the intracranial EEG recording analysis has also been experimented and compared with the bi-polar referencing scheme to make sure that the bi-polar subtraction did not remove signals. After manual inspection, we chose to use the bi-polar referencing for the further analyses.

To extract temporal-frequency information, the continuous bipolar voltage traces were decomposed using *Morlet* wavelet filtering (at log-spaced frequencies between 1 and 256 Hz [59 total frequencies]; each wavelet having a width of 5 cycles). The output instantaneous power timeseries has a sampling rate of 200 at each frequency. The timeseries was then epoched, baseline corrected and averaged across good trials.

Values in the power spectrogram were then log-transformed to approximate normal distribution and z-scored across time and frequency. To measure how EPs are consistent across trials, we calculated the inter-trial phase coherence (ITPC) by taking the complex phase component of the wavelet decomposition, averaging across trials, and then taking the absolute. Values in the ITPC spectrogram were then square-root transformed to approximate normal distribution and then z-scored across time and frequencies. The remaining text of the paper will refer to the evoked potentials (EP), the evoked power and ITPC spectrograms as their preprocessed version by default.

### Activation detection using semi-supervised Uniform Manifold Approximation and Projection (UMAP)

#### Preliminary activation labelling

The key to determining a casual connectivity lies on correctly detecting a true physiological activation among the volume conductions of an electrical stimulation. We employed a semi-supervised learning approach utilizing the Uniform Manifold Approximation and Projection (UMAP) algorithm with the python toolbox *umap-learn* (https://umap-learn.readthedocs.io/en/latest/)^54^. Unlike traditional dimensionality reduction techniques, semi-supervised UMAP allows for integration of labeled and unlabeled data, using our partial knowledge to guide the manifold structure of a true activation.

Before training, we produced a preliminary activation label based on the “consensus” of the pre-defined 3 criteria: (1) “Uniform-Cutoff”: a hard threshold of EP z-score > 7 at any time point in the [10, 600] ms time window. (2) “Channel-Adaptive”: stimulation-channel specific thresholds (SD > 2) for peak heights and peak dominances of the EPs generated from the same bipolar stimulation. EP peaks were detected using the MATLAB “findpeaks” function for all peaks (positive or negative), and the peak and prominence distributions were constructed from all the detected peaks, upon which we applied the thresholds. (3) “Phase-locking”: individual specific thresholds with a confidence level of 95% in the distribution of connectivity index scores across all stimulating-recording pairs of the same subject. The connectivity index is based on ITPC, recently created to measure causal connectivity as discussed in a recent study^11^. Apart from these criteria reported from the literature, we also imposed some criteria that should be met to be physiologically alike: the voltage trace of the EP should have smooth and fast rises in the early phase (10∼60 ms), later peaks (after 250 ms) can have lower peak heights (z-score > 5), and the signal should approximate to baseline after 800 ms. When all the criteria agree, we labelled a stimulating-recording instance as either “1” (activated) or “0” (non-activated); otherwise (∼1/3 of the cases), we labelled it as “-1” (unlabeled), which is left for the algorithm to apply learned rules and decide for us.

#### Training data preparation

We applied a semi-supervised UMAP with power and ITPC spectrograms as input data, as the spectrograms explicitly quantified the rich waveform information (components and consistency) of the EPs. We prepared the training data with the following steps. We down sampled and smoothed the spectrograms by averaging the nearest neighbors to reduce the data redundancy, save working memory for the intensive computation, and to denoise and highlight the features of interest. We used characteristic time-frequency kernels to smooth (averaging the nearest neighbors) and down-sampled the spectrograms, based on Leland McInnes k-means clustering from all stimulation-evoked potential peaks, and our prior knowledge that the first 10 ms upon a stimulation onset is mainly artifact (**Fig. S5**). Therefore, we down sampled the temporal domain into 5, 25, 20, and 10 points to cover the windows of [-30, 10] ms (artifact window), [11, 117] ms, [118, 283] ms, and [284, 800] ms. The artifact window was kept in the training data for the algorithm to learn to recognize. We used more data points for representing the [11, 283] ms because most evoked potentials happen in this time window. On the frequency domain, they were down sampled into 4 datapoints to cover the [0.5, 5] Hz (delta), [5, 8] Hz (theta), [8, 15] Hz (alpha), [15, 30] Hz (beta), [30, 70] Hz (gamma) and > 70 Hz (high). The down sampled power and ITPC spectrograms were then vectorized and concatenated as a single vector of multiple features (number of features = timepoints*frequencies of power + timepoints*frequencies of ITPC) for each stimulating-recording instance. Each stimulating-recording vector was then concatenated to be the input training data. The input data was curated using the *sklearn* toolbox in python (https://scikit-learn.org/stable/), including missing data imputation with mean statistics (negligible cases, about 0.089% of the datapoints), and quantile transformation to distribute the data normally. When both transformed and un-transformed data were experimented, the results did not change but showed a clearer clustering effect using the transformed data; hence, the results using transformed data are reported in this paper.

#### Subject-level semi-supervised UMAP learning

The number of latent dimensions for the UMAP mapping was set to 2, the size of the local neighborhood was set to 5, the minimum distance of the points in the latent space was set to 0.01. The rest of the parameters were kept at default. As a result, each participant’s EP data was classified into activation and non-activation clusters that were grouped at the far ends in latent space, while the few floating data points that were not clustered were considered outliers. Each person’s clustering was manually checked by (1) visualizing the spectrograms of the centroid and boundary points of each cluster, and (2) visualizing the inverse projection of the original space from the latent space. 26 out of 27 participants’ data were grouped into two meaningful clusters, while one participant’s data (S21_196) with extensive amount of noisy data failed to cluster and was excluded from the following analyses.

### Neural feature extraction for activated signals

#### Group-level UMAP learning

To gain an overview of the data structure of the activated spectrograms, we applied an unsupervised UMAP with input data of denoised power and ITPC spectrograms of the activated channels (number of latent dimensions = 4). To minimize the influence of electrical artifacts, we created a non-activated spectral information (power and ITPC) template for each stimulating channel by averaging all the non-activated pairs stimulating from this channel. This non-activated spectral template contains mostly the artifact of volume conduction (**Fig. S5**), as the physiological events would be averaged out when not synchronized by an activation. We used the non-activated spectral template as a control and subtracted it from all the activated spectrograms from the same stimulation channel within the same participant. When the non-activated cases for a stimulation channel are too few (< 10) and thus unable to generate a stable mean as a contrast, this happened only once, we excluded the case from the subsequent analyses. The remaining traces of the activated spectral information were considered denoised and fed into the group-level analysis.

We applied a supervised UMAP with input data of denoised power and ITPC spectrograms of the activated channels, to map to the anatomical labeling of interest. The provided anatomical labels were “THAL-ipsi”, “THAL-contr”, “COR-ipsi” and “COR-contr”, corresponding to *stimulating* from thalamus/cortex and recording from the ipsilateral/contralateral hemisphere.

The same training data preparation as mentioned before was applied. The number of latent dimensions for the UMAP mapping was set to 2, the size of the local neighborhood was set to 15, the minimum distance of the points in the latent space was set to 0.1. These parameters varied from the within-subject semi-supervised UMAP learning because the data variability within-subject is assumed to be smaller than the group data, while we aim to find commonalities that override individual differences. Therefore, the local neighborhood and the latent space are relaxed to bigger numbers (As a general principle, we experimented with different sets of parameters [within a reasonable range] to achieve the best clustering results from visual inspection). The rest of the parameters were kept at default. We further employed a hierarchical-density clustering algorithm to define the boundaries of the embedding clusters against the background noise. Finally, we investigated the centroid and averaged the presentation of the data in each cluster to verify that the clustering preserved meaningful and distinct features within the data.

To ensure the data-driven clusters are generalizable and meaningful, we fed the resulting embedding to a series of feature evaluation procedures, including using a support vector classifier to test their generalizability to the ¼ unseen testing data. We showed that the embedding for separating the cortical and thalamus evoked responses has 75.9% predictability to the unseen data, much higher than chance rate (25%). Especially, the clusters of thalamus and cortex stimulated data are mostly well preserved, suggesting these two types of connectivity show most distinctive features as encapsulated by their power and ITPC spectrograms. We further employed a hierarchical-density clustering algorithm^55^ to define the boundaries of the embedding clusters against the background noise, using the *HDBscan* toolbox (https://hdbscan.readthedocs.io/en/latest/). We then investigated the averaged spectrograms (i.e. the cluster centroid) for each cluster, for a sanity check of the clustering results.

#### Cluster-based permutation test

Category-specific spectral features were determined by cluster-based permutation significance testing: (1) one-sample T-test for determining the significant clusters for each category, and (2) paired T-test for determining the significant differences between categories (with a 2 [THAL/COR]-by-2 [ipsi/contr] design). Specifically, two-sample t-tests were conducted within subjects and the individual statistics (t-scores) were fed into a group-level one-sample T-test for final inference. The testing was performed using the toolbox MNE-python (v1.5.0) (https://mne.tools/stable/index.html). The number of permutations was set to 5000, the initial cluster forming threshold was bigger than 6 t-scores, and the cluster-wise permutation confidence interval was set to 99% (i.e., *P*_cluster_ < 0.01).

### Neural feature decoding

Based on the cluster-based permutation test, we identified three distinct neural features (i.e., significant clusters) that are specific to each anatomical category of interest (i.e., UMAP localizers). They are represented by the power and ITPC patterns in the window of ([10, 50] ms of the COR-ipsi category (averaged across all instances in the corresponding category), in the window of [70, 160] ms of the COR-contr category, and the [165, 280] in the THAL-contr category. The temporal-frequency relationships in these time ranges and anatomical categories were considered as neural features. They were decoded in each instance of stimulated/record pairs. To do this, we utilized the spectral information from the significant cluster in the group-level spectrograms as a template. We employed a sliding-window cross-correlation to match this template with the specific instances of the spectrograms, point by point in time. Different similarity (or distance) measures were experimented and turned out to produce similar results; therefore, we used the Pearson correlation *r* for its easy interpretability. The resulting timeseries of *r* has a samping rate of 200 per second, which is the same as the spectral dataset.

A sliding-window cross-correlation with the 3 identified neural features generated three temporal similarity curves for each stimulating-recoding instance, each corresponding to the neural feature’s temporal evolvement in this specific pair of stimulated/recorded sites. With the temporal similarity curve, we detected the peak similarity and the time to the peak using MATLAB “findpeak” function. Instances within each anatomical area were averaged for brain regional inferences.

### General framework of statistical testing

The datasets in the present study generally have a nested structure such that each condition consists of multiple electrode contacts, each with a brain area and subject identity; therefore, the mixed linear model was used as a general testing framework for model fitting, using the brain area and subject identifiers as nested random factors. Therefore, the comparisons were always made within the same stimulation/recording channels, brain areas and subjects, whenever possible. Practical model setups can vary slightly according to data structures and tested hypotheses. For significance inference, the model comparison approach with a hypothesis testing framework was adopted. Namely, a null model is compared with an alternative model which has the hypothesized effect added on top of the null model. The model with higher evidence is selected, based on a likelihood-ratio test. We additionally provided the Akaike information criterion (AIC) and Bayesian information criterion (BIC) in our supplementary table of post-hoc tests where spurious significant results may arise given the large numbers of tests, even after the false-discovery-rate (FDR) correction. The model fittings and selections were conducted using the *lme4* toolbox and the *anova* function in R. The model specifications were provided along with the significance tables in the supplementary materials.

## Acknowledgments

We are grateful to the patients who agreed to participate in our research, and the staff in the EEG lab including clinical fellows and attendings whose help enabled this research. We express our gratitude to our colleagues Riki Matsumoto (Kobe - Japan), Corey Keller (Stanford - USA) and Konrad Körding (Philadelphia – USA) for their valuable feedback on our findings; and to Leland McInnes (Ottawa – Canada) for running a sanity check on our UMAP analysis approach.

## Author contributions

DL contributed to the conception, acquisition, analysis, and interpretation of data and creation of new codes and measures used in the work and writing of the manuscript; JRS contributed to the analysis and interpretation of data; ZL contributed to the acquisition of data; JP contributed to conception and design of the work, acquisition and interpretation of data and writing of the manuscript. All authors provided significant feedback throughout the study and manuscript preparation and have approved the submitted version.

## Supplementary Materials

Materials and Methods

Supplementary Text

Figs. S1 to S10

Tables S1 to S8

Table S9 in a separate file (a large table with all post-hoc comparison results)

References (*##*–*##*)

Movies S1

## SUPPLEMENTARY MATERIALS

**Fig. S1.**
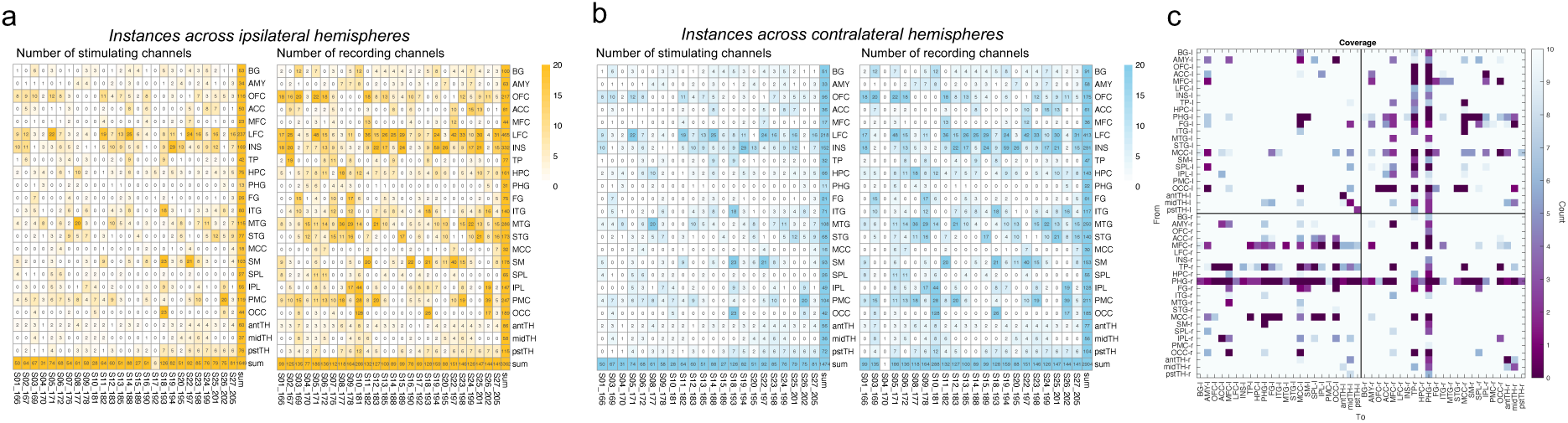
Electrode coverage. (a) and (b) list the numbers of stimulating/recording channels having been tested in each participant, across ipsilateral (yellow) and contralateral (blue) hemispheres, respectively. (d) Counts of bipolar channels stimulated and recorded across two hemispheres (l: left, r: right), highlighted at the lower coverage (n < 10). Most of the examined causal connectivity instances have a sample size >= 10.

**Fig. S2:**
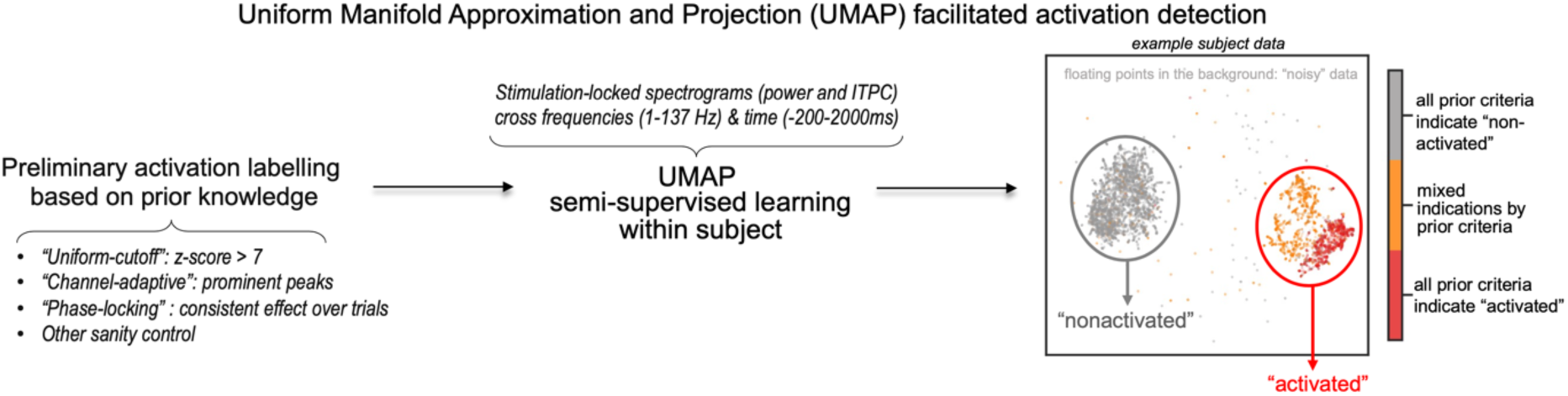
Illustration of the activation detection pipeline using semi-supervised UMAP learning. The activation detection was conducted at the individual basis. The resulting clusters have been manually checked for meaningful representation of the data.

**Fig. S3:**
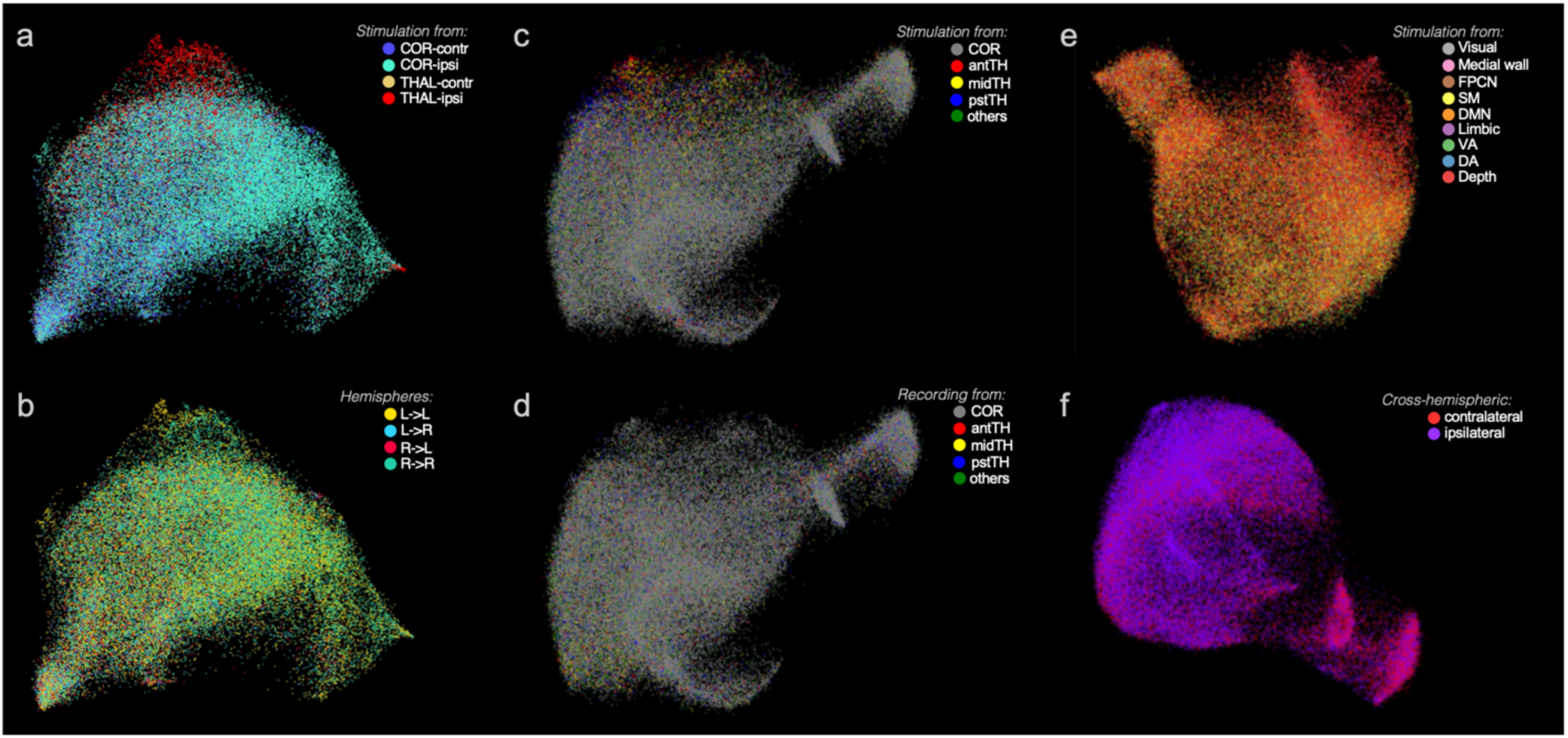
Unsupervised UMAP embedding space exploration, presented to support the anatomical category that we chose to focus on in the study, i.e., shown in (a). The same embedding structure is rotated for the best visualization. Subplots (a), (b) present the same slice of the embedding space, colored depending on (a) where the EP is originated from (regardless of the recording site’s location), and (b) which hemisphere the SEP is stimulated and recorded from. Subplots (c) and (d) present the same slice of the embedding space, but they differ in their color correspondence to the stimulation sites’ location (COR: cortex, antTH: anterior thalamus, midTH: mid thalamus, pstTH: posterior thalamus, others: other subcortical areas including amygdala, basal ganglia, claustrum). Subplots (e) and (f) show the same embedding space, from different viewing angles, colored to (e) the large-scale intrinsic network identities of the stimulation sites, and (f) the cross-hemispheric condition of a SEP (i.e., ipsilateral SEPs are the pairs stimulated and recorded from the same hemisphere, regardless of the left or right hemisphere; and the contralateral SEPs are those stimulated and recorded from the other hemisphere) (DMN: default-mode network, FPCN: fronto-parietal control network, SM: sensori-motor network, VA: ventral attention network, DA: dorsal attention network). The examples show that the recording sites’ location, intrinsic network territories and hemisphere identity are not determining factors of the SEP’s spectral profiles; rather, whether the connectivity initiates from the thalamus or cortex andspreads to the ipsilateral or contralateral hemisphere, produces distinct spectral patterns of the EP.

**Fig. S4:**
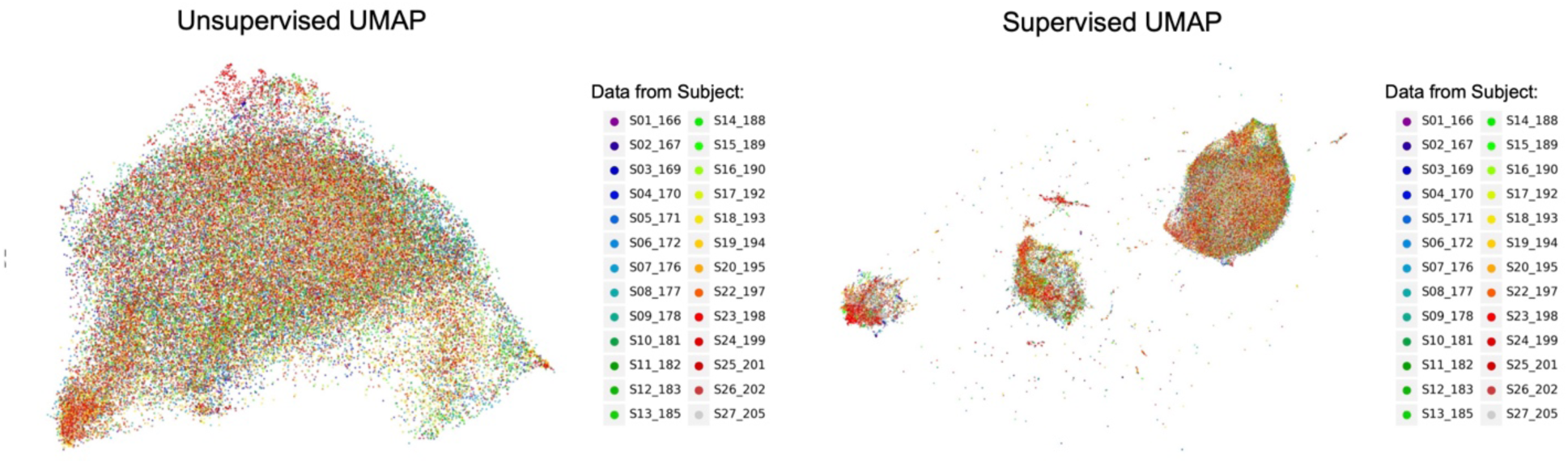
Mapping between the UMAP embedding spaces with subject identity (with S21 excluded from the analyses due to excessive number of bad channels in this subject’s data). The unsupervised and supervised UMAP embedding spaces were the same as presented in the main article, but colored with subject ID, instead of anatomical labelling. The subject-ID distributions do not correspond to the embedding structures, suggesting that the embedding spaces and clustering are not dominated by the individual differences which may be suspected as a confounding factor.

**Fig. S5.**
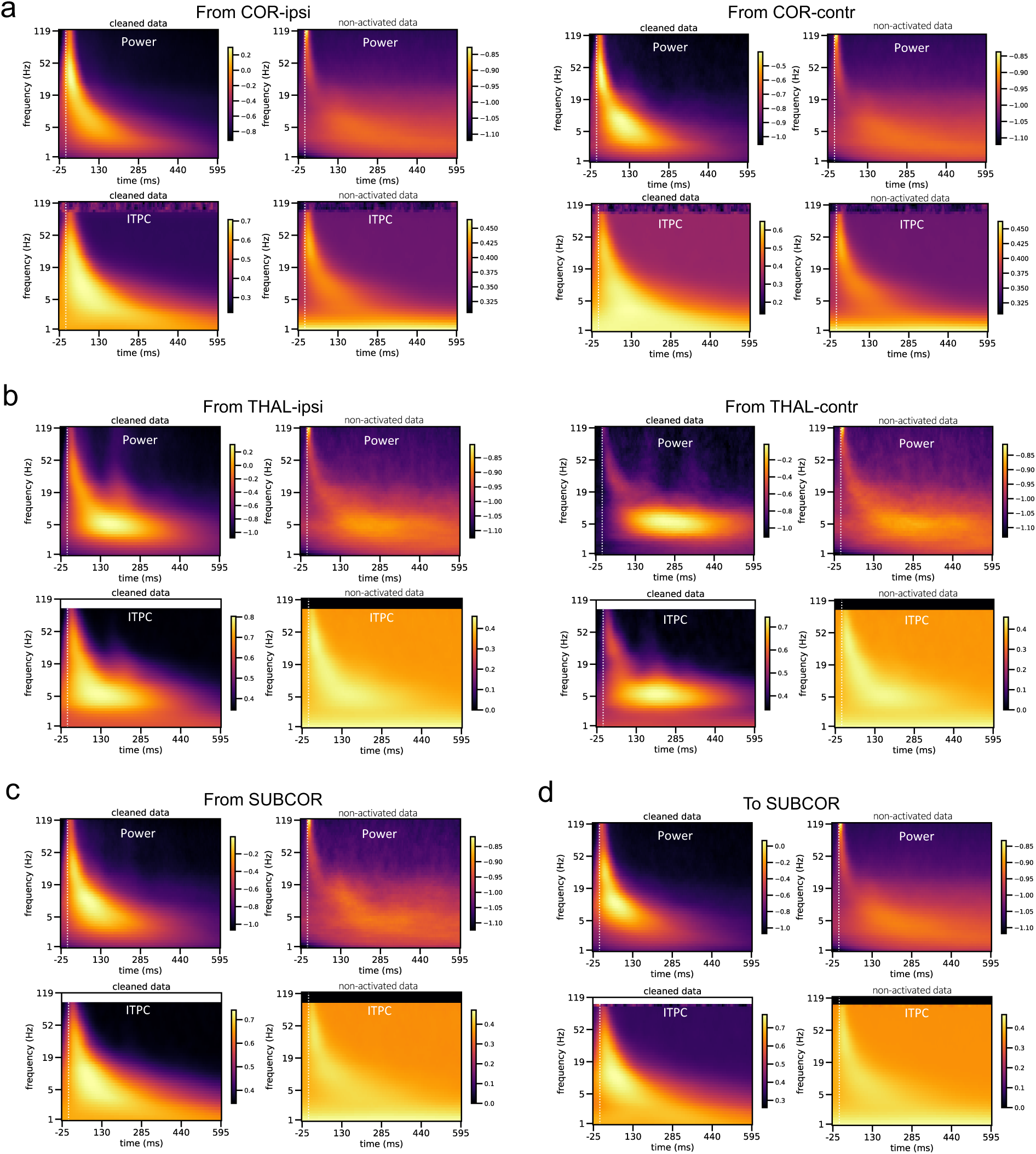
Group-averaged power and ITPC, cleaned data (left) and noisy non-activated data (right) that is subtracted from the channel-matched spectrograms by which the cleaned data was obtained. To minimize the influence of electrical artifacts, we created a non-activated spectral information (power and ITPC) template for each stimulating channel by averaging all the non-activated pairs stimulating from this channel. This non-activated spectral template contains mostly the artifact of volume conduction, hallmarked by the high-frequency peak pixel immediately following the stimulation onset. We used the non-activated spectral template as a control and subtracted it from all the activated spectrograms from the same stimulation channel within the same participant. The remaining traces of the activated spectral information were considered cleaned data. The colorbars indicate the magnitudes for power and ITPC, which are log-transformed and z-scored for power and square-root transformed and z-scored for ITPC. (a) Spectrograms averaged across all instances with stimulating channels from the ipsilateral (COR-ipsi) or contralateral (COR-contr) cortices. (b) Spectrograms averaged across all instances with stimulating channel from the ipsilateral (THAL-ipsi) or contralateral (THAL-contr) thalamus. (c) and (d) show spectrograms averaged across all instances with stimulating or recording channels, respectively, in the non-thalamic subcortical areas. We note that the F1 feature cannot be an artifact of volume conduction because i) electrical artifact appears to be a highlighted pixel at the high frequency (>100Hz) and immediately upon the stimulation (within 10 ms) on the power spectrogram (see non-activated data). It has a very different characteristic than F1. ii) All spectrograms for activated data have been subtracted with the non-activated data (containing the stimulation artifact feature) before statistical testing. iii) In terms of the feature decoding for individual connections, we relied on multi-variate pattern matching in the time-frequency domain which is not biased by the amplitude in the time domain.

**Fig. S6:**
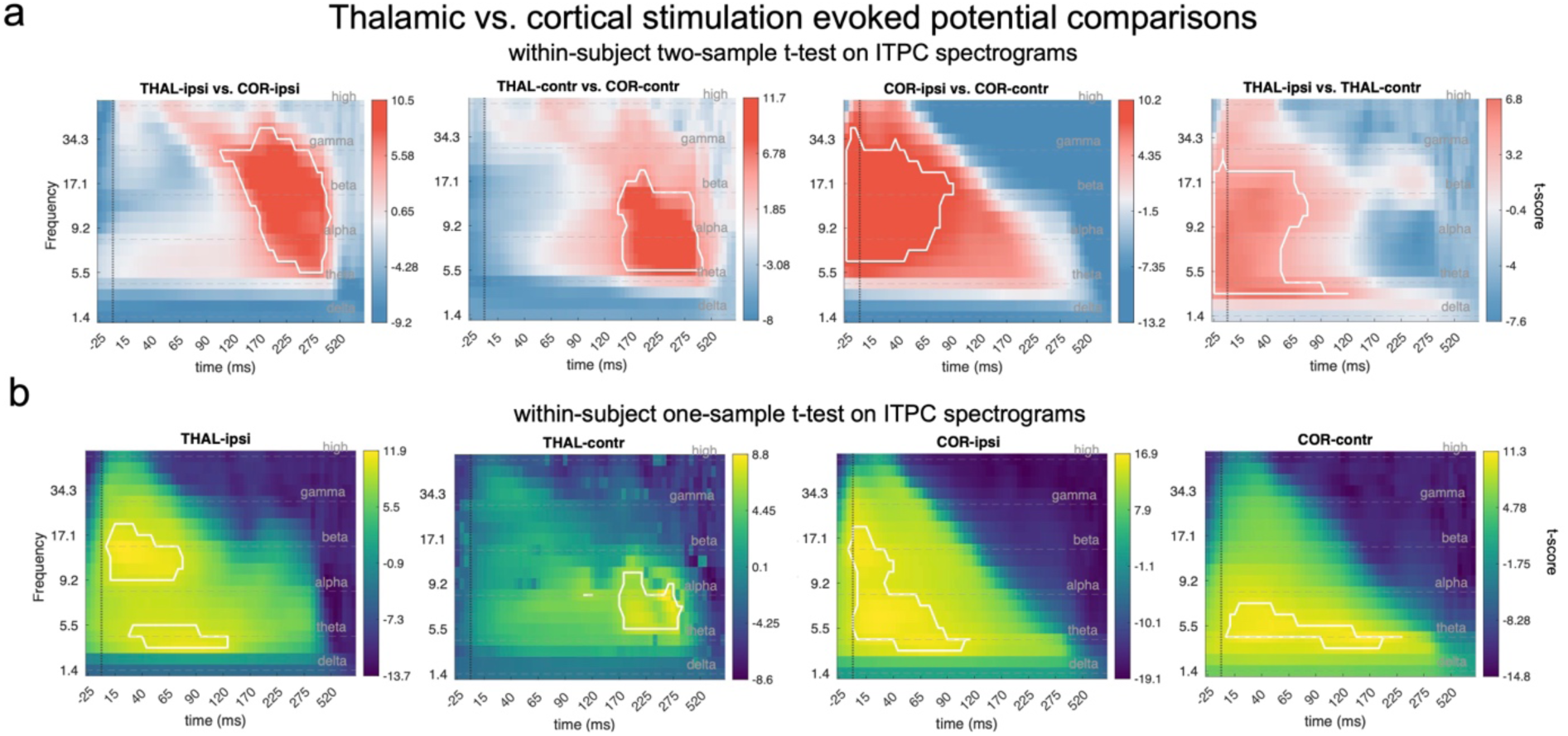
Spectrograms of stimulation-evoked inter-trial phase coherence (ITPC) within and between anatomical categories (COR: cortex, THAL: thalamus, ipsi: ipsilateral, contr: contralateral, →: causal influence direction). Highlighted contours on the spectrograms indicate significant clusters (n-permutation = 5000, initial cluster forming threshold = 6 *t*-scores, *P*_cluster_ < 0.01). Dashed lines on the spectrograms denotes the segments of conventional frequency bands: [0.5, 5] Hz (delta), [5, 8] Hz (theta), [8, 15] Hz (alpha), [15, 30] Hz (beta), [30, 70] Hz (gamma) and > 70 Hz (high). (a) Thalamic vs. cortical evoked spectrogram comparison with within-subject two-sample t-tests (two-tailed) on the ITPC spectrograms, significance inference performed with cluster-based permutation testing. White contours highlight significant clusters of the contrast indicated on the subtitle. Reversed contrasts did not generate significant results. (b) Thalamic and cortical evoked ITPC spectrograms and significant clusters in either ipsilateral or contralateral hemispheres. Significance testing conducted with within-subject one-sample t-tests. Acronym example “THAL-ipsi” on the subtitle means recording from pairs of electrodes that were stimulated from the ipsilateral thalamus.

**Fig. S7:**
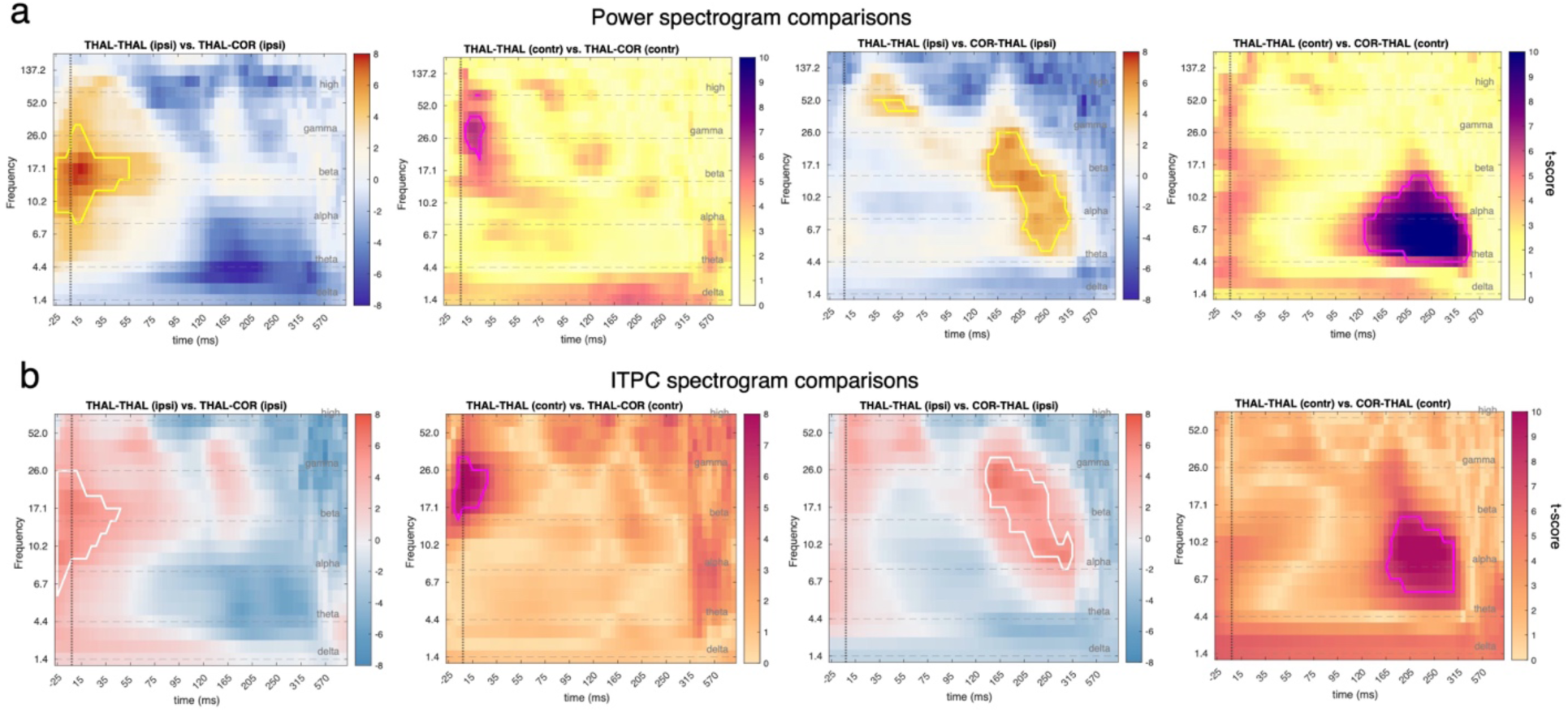
Cluster-based permutation tests among different anatomical categories of thalamus-related connectivity. (a) Power spectrogram comparisons. (b) ITPC spectrogram comparisons. Tested contrasts are between THAL→ THAL vs. THAL→COR, and between THAL→THAL vs. COR→THAL, in either ipsilateral or contralateral pairs. The tests were conducted within-subject by default, but the number of subjects with comparable contralateral THAL→ THAL instances were too few (<10) to generate reliable significance testing; therefore, the tests involving this condition were conducted with a between-subject design (i.e., two-sample t-test at the group level in the second and the fourth columns).

**Fig. S8.**
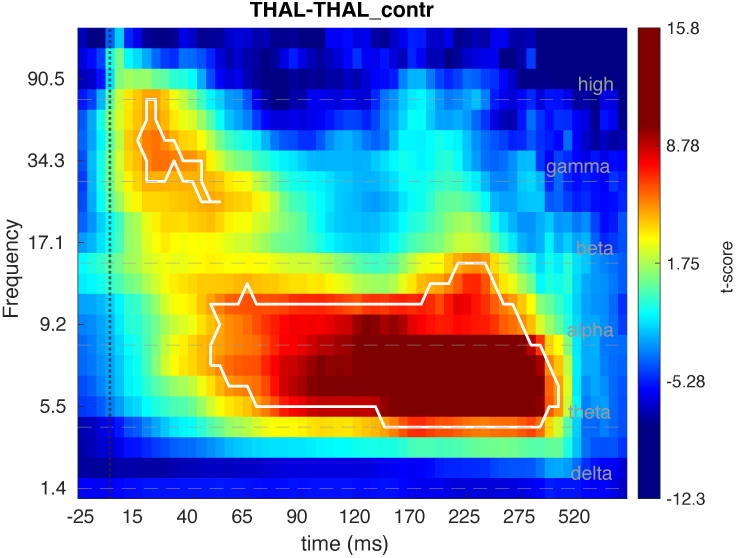
Cluster-based permutation test on THAL->THAL (contra) connections (no mixed design). The test at the individual level could not be reliably estimated due to insufficient number of subjects (n-subject = 7). However, among the 7 individuals with bilateral thalamic stimulations and recordings, there was a sufficient sample size of electrode contacts (n-connections = 81) to estimate the effect at the group level. The color-bar represents a group-level t-score of a one-sample t-test. Significance clusters are marked by white contours.

**Fig. S9.**
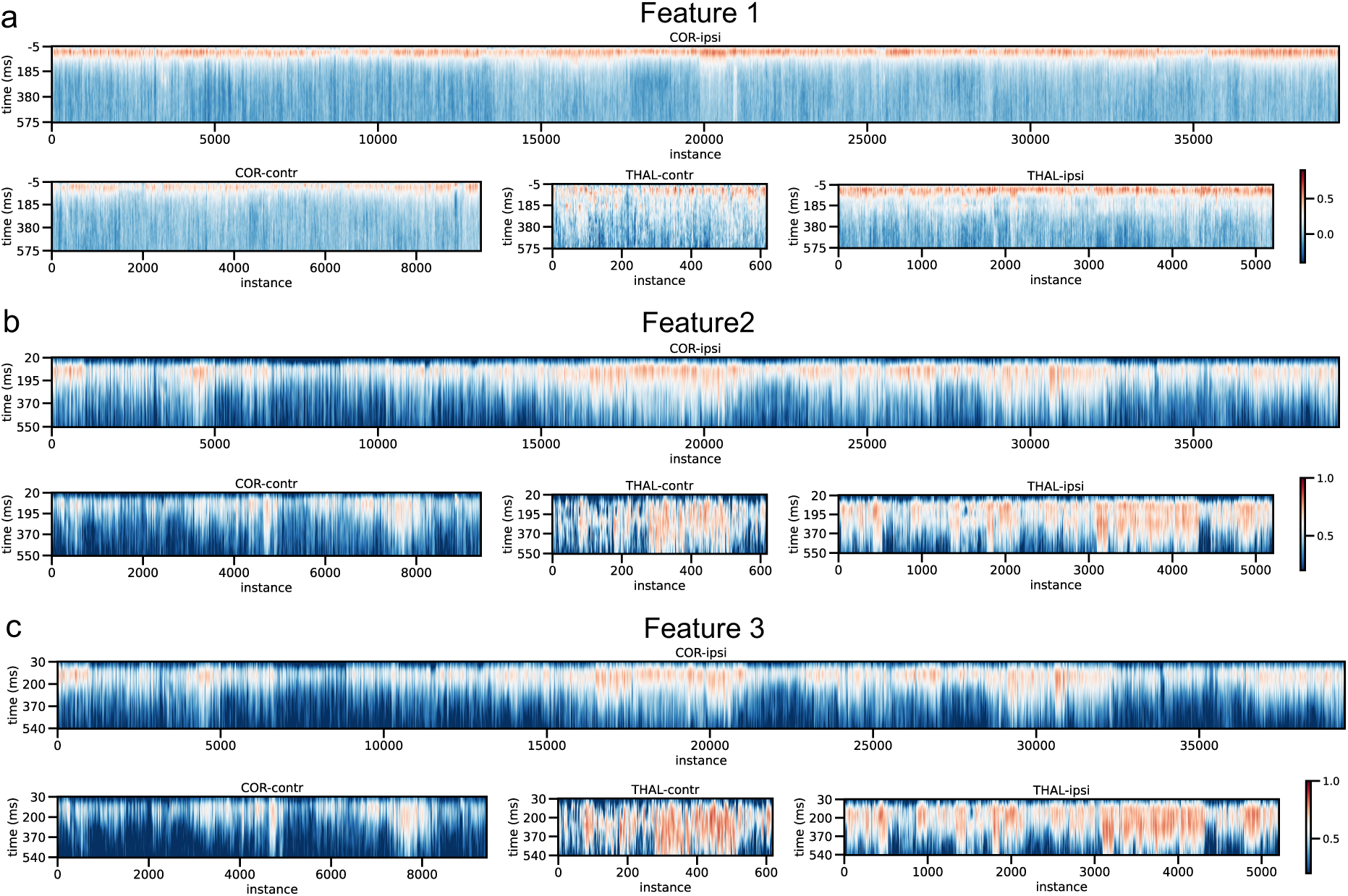
Data overview of the feature representations for each anatomical category of the stimulation channel. (a)-(c) respectively show the F1-3 representations. For each heatmap, the x-axis shows the instance number of the whole cohort. An instance is a causal-connectivity case (with a complex spectral profile depicted by the power and ITPC spectrogram) that is established by a unique stimulation-recording channel pair. Instances with the same stimulation category are grouped here, including COR-ipsi, COR-contr, THAL-ipsi and THAL-contr. The y-axis shows the time upon stimulation onset. F2 and 3 have later starting times because their spectra-temporal window is long which renders an equally long window to estimate its representation at each timepoint, while the datapoints in the beginning of the trial (including the baseline) are not sufficient for an estimation of these features. However, the F2 and 3 have a late onset which permits this limitation to be irrelevant. The color bars show the Pearson-correlation coefficient. The hotter color indicates the rising of the feature.

**Fig. S10:**
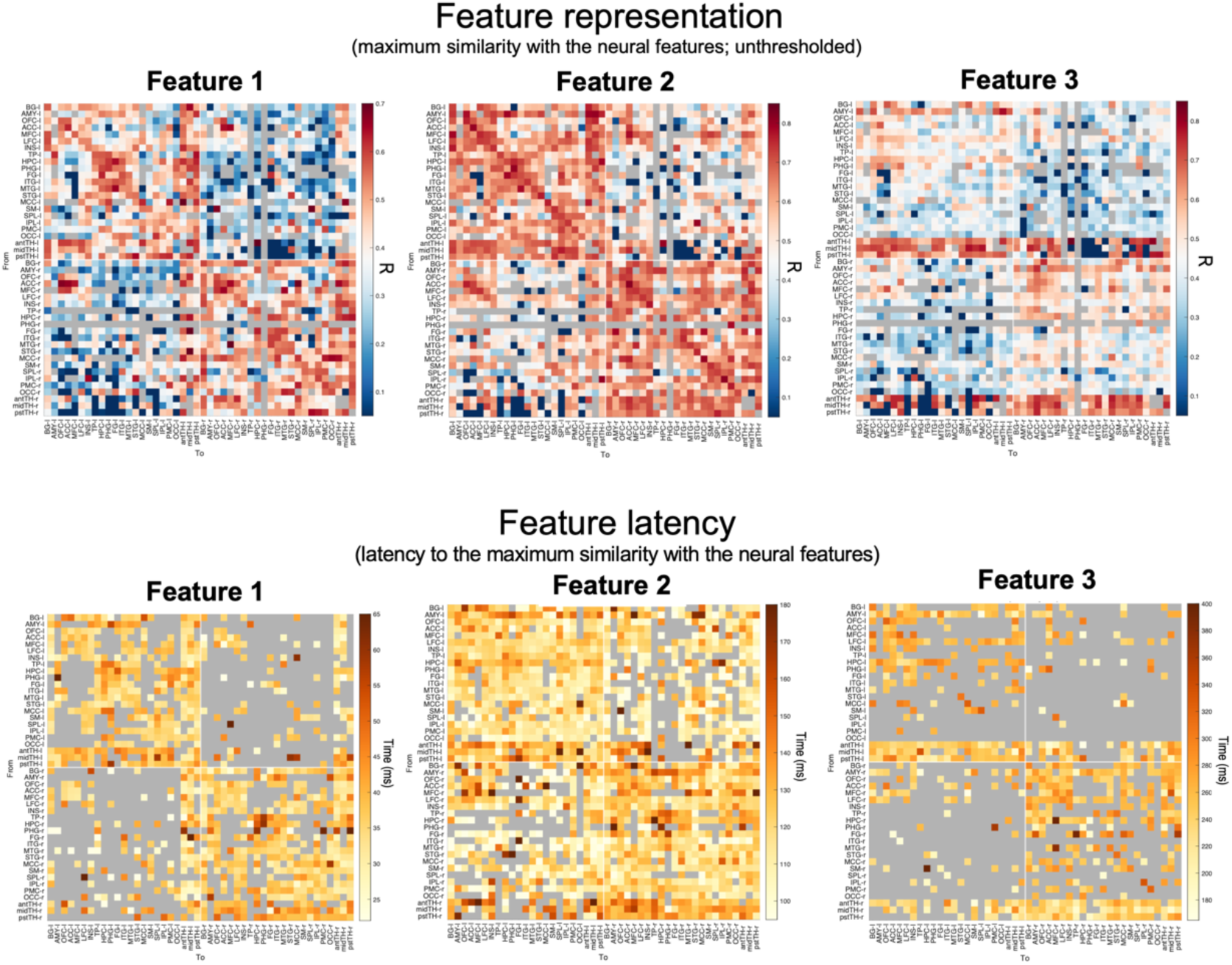
Feature representation and latency for the three neural features. For R matrices, the grey entries indicate NaN values where the sample size for this connectivity is not bigger than 10 (Fig. S1c). Otherwise, the r* (maximum similarity reached during the feature window) of all incidences in this pair of anatomical regions are averaged: 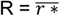. Feature latency is the time duration from the stimulation onset to the timepoint where the r* is reached. Instances with small r* (< 0.4 for Feature 1, 2 and < 0.5 for Feature 3) are neglected and counted as NaN in the grey entries in the latency matrices. Feature 2&3 generally have stronger r* because the two features persist longer and are concentrated at the lower frequencies: not only do longer time windows provide more timepoints to correlation which provides larger statistical power, but also the lower frequencies usually have bigger magnitude on the spectrograms. Both factors contribute to the higher correlation coefficients for the Feature 2&3. To address that, we did not directly compare the R values unless normalized. We also defined the cutoffs of the R values adaptively for each feature by visually inspecting the histograms of their distributions as well as the concerned anatomical patterns.

## Supplementary Tables

**Table S1.** Evaluation of the three preliminary labelling algorithms which are based on traditional methods, with the semi-supervised UMAP clustering as a reference. Peak detection at the temporal domain using a uniform threshold (i.e., “Uniform-Cutoff”) to determine a high amplitude is mostly often used in the previous literature. The results show that it has high precision but low sensitivity, i.e., tends to make Type-II errors. Stimulation and recording of non-neocortex such as subcortex and limbic areas often evoke potentials with relatively small peaks and unconventional waveforms, in which cases traditional peak-detection methods tend to fail.

| *Activation Criteria* | *Accuracy* | *Sensitivity* | *Precision* | *False Positive Rate* | *Correlation with UMAP learning* |
| --- | --- | --- | --- | --- | --- |
| *Uniform-Cutoff* | 0.80 | 0.37 | 0.99 | 0.01 | 0.52 |
| *Channel-Adaptive* | 0.84 | 0.53 | 0.95 | 0.01 | 0.63 |
| *Phase-Locking* | 0.91 | 0.77 | 0.99 | 0.02 | 0.80 |

### ROI analysis

#### Between-thalamic comparisons

**Table S2.** Comparison of F1 representations in the thalamic inflow pathways between antTH and pstTH. Here we tested all brain areas (of both hemispheres) that had sufficient electrode coverage, i.e., more than two electrodes covered in the area within-subject, and ten observations in total. The tested statistical model is a mixed linear model, with R as the dependent variable, predicated by (i) the fixed effect of “ROI” (i.e., pstTH minus antTH) and (ii) the random effect of the variability among the electrode contacts (i.e., the different electrode contacts to be stimulated, which are shared between the ROIs; similar to a paired t-test model). Using the toolbox of “lme4” ^56^ (https://cran.r-project.org/web/packages/lme4/), the model and the compared null model are written as the following: ml: r ∼ ROI+ (1|elecID); ml0: r ∼ (1|elecID).

| Area | NO.<br>Observations | NO.<br>Random Groups (adjusted) | Estimate | t-value | p | p (FDR-corr.) |
| --- | --- | --- | --- | --- | --- | --- |
| BG | 173 | elecID:34 | -0.08 | -2.61 | 0.01 | 0.03 |
| AMY | 138 | elecID:24 | 0.01 | 0.47 | 0.64 | 0.70 |
| OFC | 153 | elecID:42 | -0.13 | -4.74 | <<0.01 | <<0.01 |
| ACC | 99 | elecID:20 | -0.18 | -5.79 | <<0.01 | <<0.01 |
| MFC | 60 | elecID:11 | -0.15 | -3.36 | 0.00 | 0.01 |
| LFC | 446 | elecID:98 | -0.09 | -5.41 | <<0.01 | <<0.01 |
| INS | 446 | elecID:97 | -0.02 | -0.82 | 0.41 | 0.58 |
| TP | 56 | elecID:13 | -0.02 | -0.51 | 0.60 | 0.70 |
| HPC | 221 | elecID:42 | 0.03 | 1.42 | 0.16 | 0.28 |
| PHG | 35 | elecID:7 | 0.09 | 1.82 | 0.07 | 0.16 |
| FG | 46 | elecID:10 | 0.02 | 0.37 | 0.71 | 0.74 |
| ITG | 114 | elecID:28 | 0.06 | 1.81 | 0.07 | 0.16 |
| MTG | 228 | elecID:46 | 0.06 | 2.49 | 0.01 | 0.04 |
| STG | 165 | elecID:38 | -0.02 | -0.64 | 0.52 | 0.65 |
| MCC | 77 | elecID:12 | -0.10 | -2.62 | 0.01 | 0.03 |
| SM | 227 | elecID:51 | 0.05 | 2.43 | 0.02 | 0.04 |
| SPL | 22 | elecID:5 | 0.05 | 0.61 | 0.52 | 0.65 |
| IPL | 103 | elecID:19 | 0.04 | 1.20 | 0.26 | 0.40 |
| PMC | 286 | elecID:55 | 0.02 | 1.12 | 0.27 | 0.40 |
| OCC | 64 | elecID:15 | 0.06 | 1.63 | 0.10 | 0.19 |

<<: far smaller than; Estimate "positive": pstTH > antTH

**Table S3.** Comparison of F1 representations in the thalamic outflow pathways between antTH and pstTH. Again tested for all the brain areas (of both hemispheres) that have sufficient electrode coverage: more than two electrodes covered in this area within-subject, and ten observations in total. The tested statistical model is a mixed linear model, with R as the dependent variable, predicated by (i) the fixed effect of “ROI” (i.e., pstTH minus antTH) and (ii) the random effect of the variability among the electrode contacts (i.e., the different electrode contacts to be recorded from, which are shared between the ROIs; similar to a paired t-test model). Using the toolbox of “lme4” ^56^ (https://cran.r-project.org/web/packages/lme4/), the model and the compared null model are written as the following: ml: r ∼ ROI+ (1|elecID); ml0: r ∼ (1|elecID).

| Area | NO.<br>Observations | NO.<br>Random Groups (adjusted) | Estimate | t-value | p | p (FDR-corr.) |
| --- | --- | --- | --- | --- | --- | --- |
| BG | 258 | elecID:57 | -0.04 | -1.54 | 0.12 | 0.17 |
| AMY | 34 | elecID:10 | -0.07 | -1.18 | 0.25 | 0.29 |
| OFC | 102 | elecID:35 | -0.17 | -4.60 | <<0.01 | <<0.01 |
| ACC | 65 | elecID:17 | -0.24 | -6.91 | <<0.01 | <<0.01 |
| MFC | 14 | elecID:6 | -0.05 | -0.34 | 0.71 | 0.75 |
| LFC | 446 | elecID:130 | -0.13 | -7.91 | <<0.01 | <<0.01 |
| INS | 299 | elecID:85 | 0.08 | 4.04 | <<0.01 | 0.00 |
| HPC | 53 | elecID:19 | -0.03 | -0.65 | 0.51 | 0.57 |
| FG | 19 | elecID:6 | -0.09 | -1.37 | 0.16 | 0.20 |
| ITG | 62 | elecID:22 | 0.06 | 1.58 | 0.12 | 0.17 |
| MTG | 200 | elecID:71 | 0.14 | 6.59 | <<0.01 | <<0.01 |
| STG | 173 | elecID:50 | 0.22 | 9.63 | <<0.01 | <<0.01 |
| MCC | 56 | elecID:18 | -0.10 | -2.50 | 0.01 | 0.02 |
| SM | 155 | elecID:44 | 0.06 | 2.19 | 0.03 | 0.05 |
| SPL | 25 | elecID:9 | 0.28 | 3.57 | 0.00 | 0.01 |
| IPL | 159 | elecID:41 | 0.20 | 9.26 | <<0.01 | <<0.01 |
| PMC | 308 | elecID:83 | 0.18 | 9.46 | <<0.01 | <<0.01 |
| OCC | 19 | elecID:8 | 0.27 | 3.14 | 0.00 | 0.01 |
| midTH | 30 | elecID:12 | -0.01 | -0.17 | 0.87 | 0.87 |

<<: far smaller than; Estimate "negative": pstTH > antTH

**Table S4.** Comparison of F3 representations in the thalamic inflow pathways between antTH and pstTH. Again tested for all the brain areas (of both hemispheres) that have sufficient electrode coverage: more than two electrodes covered in this area within-subject, and ten observations in total. The tested statistical model is a mixed linear model, with R as the dependent variable, predicated by (i) the fixed effect of “ROI” (i.e., pstTH minus antTH) and (ii) the random effect of the variability among the electrode contacts (i.e., the different electrode contacts to be stimulated, which are shared between the ROIs; similar to a paired t-test model). Using the toolbox of “lme4” ^56^ (https://cran.r-project.org/web/packages/lme4/), the model and the compared null model are written as the following: ml: r ∼ ROI+ (1|elecID); ml0: r ∼ (1|elecID).None of the comparisons were significant: it is expectable because F3 is not common in the thalamic inflow pathways, hence the comparison of cortex-evoked F3 signals in the thalamus between antTH and pstTH only show spurious differences.

| Area | NO.<br>Observations | NO.<br>Random Groups (adjusted) | Estimate | t-value | p | p (FDR-corr.) |
| --- | --- | --- | --- | --- | --- | --- |
| BG | 173 | elecID:34 | -0.08 | -2.61 | 0.01 | 0.10 |
| AMY | 138 | elecID:24 | -0.01 | -0.28 | 0.79 | 0.87 |
| OFC | 153 | elecID:42 | -0.07 | -2.46 | 0.01 | 0.10 |
| ACC | 99 | elecID:20 | -0.08 | -2.50 | 0.01 | 0.10 |
| MFC | 60 | elecID:11 | 0.02 | 0.41 | 0.69 | 0.81 |
| LFC | 446 | elecID:98 | -0.04 | -2.15 | 0.03 | 0.17 |
| INS | 446 | elecID:97 | -0.04 | -2.01 | 0.04 | 0.19 |
| TP | 56 | elecID:13 | 0.04 | 0.84 | 0.40 | 0.61 |
| HPC | 221 | elecID:42 | 0.03 | 0.89 | 0.38 | 0.61 |
| PHG | 35 | elecID:7 | 0.04 | 0.64 | 0.53 | 0.69 |
| FG | 46 | elecID:10 | 0.01 | 0.16 | 0.88 | 0.92 |
| ITG | 114 | elecID:28 | 0.04 | 1.09 | 0.27 | 0.61 |
| MTG | 228 | elecID:46 | 0.02 | 0.77 | 0.44 | 0.61 |
| STG | 165 | elecID:38 | -0.02 | -0.57 | 0.56 | 0.69 |
| MCC | 77 | elecID:12 | -0.06 | -1.26 | 0.21 | 0.60 |
| SM | 227 | elecID:51 | -0.03 | -1.21 | 0.23 | 0.60 |
| SPL | 22 | elecID:5 | -0.09 | -0.89 | 0.35 | 0.61 |
| IPL | 103 | elecID:19 | 0.06 | 1.27 | 0.21 | 0.60 |
| PMC | 286 | elecID:55 | -0.02 | -0.82 | 0.41 | 0.61 |
| OCC | 64 | elecID:15 | 0.00 | 0.03 | 0.97 | 0.97 |
| midTH | 33 | elecID:11 | 0.03 | 0.86 | 0.38 | 0.61 |

Estimate "positive": *pstTH* > *antTH*

**Table S5.** Comparison of F3 representations in the thalamic outflow pathways between antTH and pstTH. Again tested for all the brain areas (of both hemispheres) that have sufficient electrode coverage: more than two electrodes covered in this area within-subject, and ten observations in total. The tested statistical model is a mixed linear model, with R as the dependent variable, predicated by (i) the fixed effect of “ROI” (i.e., pstTH minus antTH) and (ii) the random effect of the variability among the electrode contacts (i.e., the different electrode contacts to be recorded from, which are shared between the ROIs; similar to a paired t-test model). Using the toolbox of “lme4” ^56^ (https://cran.r-project.org/web/packages/lme4/), the model and the compared null model are written as the following: ml: r ∼ ROI+ (1|elecID); ml0: r ∼ (1|elecID).

| Area | NO.<br>Observations | NO.<br>Random Groups (adjusted) | Estimate | t-value | p | p (FDR-corr.) |
| --- | --- | --- | --- | --- | --- | --- |
| BG | 258 | elecID:57 | -0.07 | -2.85 | 0.00 | 0.02 |
| AMY | 34 | elecID:10 | -0.05 | -0.72 | 0.47 | 0.64 |
| OFC | 102 | elecID:35 | -0.12 | -3.12 | 0.00 | 0.02 |
| ACC | 65 | elecID:17 | -0.06 | -2.31 | 0.03 | 0.08 |
| MFC | 14 | elecID:6 | -0.14 | -2.05 | 0.07 | 0.12 |
| LFC | 446 | elecID:130 | -0.11 | -7.53 | <<0.01 | <<0.01 |
| INS | 299 | elecID:85 | -0.05 | -2.21 | 0.03 | 0.08 |
| HPC | 53 | elecID:19 | -0.11 | -2.47 | 0.02 | 0.07 |
| FG | 19 | elecID:6 | -0.15 | -2.00 | 0.05 | 0.10 |
| ITG | 62 | elecID:22 | -0.03 | -0.66 | 0.50 | 0.64 |
| MTG | 200 | elecID:71 | -0.06 | -3.08 | 0.00 | 0.02 |
| STG | 173 | elecID:50 | 0.00 | -0.09 | 0.93 | 0.97 |
| MCC | 56 | elecID:18 | -0.08 | -2.02 | 0.05 | 0.10 |
| SM | 155 | elecID:44 | -0.04 | -1.15 | 0.25 | 0.37 |
| SPL | 25 | elecID:9 | 0.03 | 0.41 | 0.68 | 0.81 |
| IPL | 159 | elecID:41 | 0.00 | 0.04 | 0.97 | 0.97 |
| PMC | 308 | elecID:83 | -0.03 | -1.93 | 0.05 | 0.10 |
| OCC | 19 | elecID:8 | -0.11 | -1.15 | 0.25 | 0.37 |
| midTH | 30 | elecID:12 | -0.01 | -0.19 | 0.84 | 0.94 |

<<: far smaller than; Estimate "positive": pstTH > antTH

**Table S6.** Significance testing results for the comparison of late thalamocortical (F3-outflow) > early thalamocortical (F1-outflow) in different brain areas. The tests were conducted only for ipsilateral pairs (i.e., regional contacts from the same hemisphere of the thalamic contacts, regardless of the left or right hemispheres), because F1 is not commonly present in the contralateral outflow pathway of the thalamus. The tests used a mixed linear model, with the z-scored r* as the dependent variable, predicated by (i) the fixed effect of *pathway* (i.e., “F3-outflow” vs. “F1-outflow”), (ii) the random effect of the coupled-contacts identity for a connectivity pair (i.e., the repeated measure of the electrode contacts concerned in both inflow and outflow pathways), (iii) the random effect of the thalamic sites being considered (ii) the random effect of individual differences. Model comparison is conducted to test the significance of the fixed effect of *pathway*. Using the toolbox of “lme4” ^56^ (https://cran.r-project.org/web/packages/lme4/), the model and the compared null model are written as the following: ml: zr ∼ pathway + (1 | pairID) + (1 | seed_elecID) + (1 | subject); ml0: zr ∼ (1 | pairID) + (1 | seed_elecID) + (1 | subject).

| ROI | Area | NO.<br>Observations | NO.<br>Random Groups (adjusted) | Estimate | t-value | p | p (FDR-corr.) |
| --- | --- | --- | --- | --- | --- | --- | --- |
| antTH | BG | 320 | pairID:153, seed_elecID:48, subject:19 | 0.12 | 1.35 | 0.18 | 0.25 |
| antTH | AMY | 68 | pairID:34, seed_elecID:23, subject:11 | 0.7 | 3.58 | <<0.01 | <<0.01 |
| antTH | OFC | 464 | pairID:228, seed_elecID:40, subject:17 | 0.04 | 0.61 | 0.54 | 0.57 |
| antTH | ACC | 226 | pairID:113, seed_elecID:25, subject:10 | 0.18 | 1.99 | 0.05 | 0.08 |
| antTH | MFC | 66 | pairID:33, seed_elecID:13, subject:7 | 0.45 | 2.21 | 0.03 | 0.05 |
| antTH | LFC | 1102 | pairID:550, seed_elecID:50, subject:22 | 0.2 | 4.22 | <<0.01 | <<0.01 |
| antTH | INS | 498 | pairID:245, seed_elecID:53, subject:24 | 0.61 | 7.58 | <<0.01 | <<0.01 |
| antTH | TP | 74 | pairID:35, seed_elecID:9, subject:7 | 0.83 | 4.74 | <<0.01 | <<0.01 |
| antTH | HPC | 132 | pairID:62, seed_elecID:30, subject:18 | 0.24 | 1.63 | 0.1 | 0.15 |
| antTH | PHG | 28 | pairID:14, seed_elecID:8, subject:6 | 0.51 | 2.34 | 0.03 | 0.05 |
| antTH | FG | 54 | pairID:26, seed_elecID:13, subject:8 | 0.31 | 1.74 | 0.08 | 0.13 |
| antTH | ITG | 150 | pairID:75, seed_elecID:22, subject:14 | 0.58 | 4.83 | <<0.01 | <<0.01 |
| antTH | MTG | 416 | pairID:201, seed_elecID:38, subject:20 | 1.13 | 16 | <<0.01 | <<0.01 |
| antTH | STG | 182 | pairID:89, seed_elecID:29, subject:14 | 0.39 | 2.73 | 0.01 | 0.01 |
| antTH | MCC | 76 | pairID:37, seed_elecID:11, subject:6 | 0.18 | 1.12 | 0.26 | 0.31 |
| antTH | SM | 184 | pairID:92, seed_elecID:25, subject:14 | 0.63 | 4.88 | <<0.01 | <<0.01 |
| antTH | SPL | 46 | pairID:23, seed_elecID:10, subject:8 | 0.75 | 3.88 | <<0.01 | <<0.01 |
| antTH | IPL | 176 | pairID:86, seed_elecID:21, subject:10 | 0.91 | 8.2 | <<0.01 | <<0.01 |
| antTH | PMC | 390 | pairID:191, seed_elecID:35, subject:17 | 1.02 | 12.93 | <<0.01 | <<0.01 |
| antTH | OCC | 56 | pairID:28, seed_elecID:7, subject:5 | 0.95 | 4.97 | <<0.01 | <<0.01 |
| antTH | antTH | 18 | pairID:9, seed_elecID:8, subject:5 | 0.88 | 3.72 | <<0.01 | 0.01 |
| antTH | midTH | 48 | pairID:24, seed_elecID:13, subject:10 | 0.03 | 0.16 | 0.87 | 0.87 |
| antTH | pstTH | 158 | pairID:75, seed_elecID:34, subject:17 | 0.16 | 1.25 | 0.21 | 0.28 |
| midTH | BG | 238 | pairID:119, seed_elecID:28, subject:10 | 0.08 | 0.73 | 0.46 | 0.51 |
| midTH | AMY | 56 | pairID:28, seed_elecID:18, subject:6 | 0.8 | 4.83 | <<0.01 | <<0.01 |
| midTH | OFC | 240 | pairID:120, seed_elecID:23, subject:9 | 0.54 | 5.22 | <<0.01 | <<0.01 |
| midTH | ACC | 152 | pairID:76, seed_elecID:14, subject:6 | 0.88 | 6.96 | <<0.01 | <<0.01 |
| midTH | MFC | 80 | pairID:40, seed_elecID:12, subject:6 | 0.33 | 1.99 | 0.05 | 0.08 |
| midTH | LFC | 716 | pairID:358, seed_elecID:33, subject:12 | 0.59 | 11.14 | <<0.01 | <<0.01 |
| midTH | INS | 334 | pairID:167, seed_elecID:33, subject:13 | 0.87 | 9.31 | <<0.01 | <<0.01 |
| midTH | TP | 8 | pairID:4, seed_elecID:3, subject:2 | 1.57 | 9.07 | <<0.01 | <<0.01 |
| midTH | HPC | 50 | pairID:25, seed_elecID:15, subject:7 | 1.01 | 5.31 | <<0.01 | <<0.01 |
| midTH | PHG | 2 | NA | NA | NA | NA | NA |
| midTH | FG | 14 | pairID:7, seed_elecID:3, subject:3 | -0.25 | -0.47 | 0.62 | 0.63 |
| midTH | ITG | 82 | pairID:41, seed_elecID:13, subject:5 | 0.65 | 4.24 | <<0.01 | <<0.01 |
| midTH | MTG | 192 | pairID:96, seed_elecID:24, subject:10 | 1.32 | 13.62 | <<0.01 | <<0.01 |
| midTH | STG | 64 | pairID:32, seed_elecID:16, subject:10 | 0.23 | 1.05 | 0.29 | 0.34 |
| midTH | MCC | 46 | pairID:23, seed_elecID:10, subject:5 | 0.2 | 1.17 | 0.24 | 0.3 |
| midTH | SM | 278 | pairID:139, seed_elecID:19, subject:6 | 0.82 | 7.94 | <<0.01 | <<0.01 |
| midTH | SPL | 20 | pairID:10, seed_elecID:5, subject:3 | 0.94 | 3.37 | <<0.01 | 0.01 |
| midTH | IPL | 116 | pairID:58, seed_elecID:14, subject:7 | 1.26 | 10 | <<0.01 | <<0.01 |
| midTH | PMC | 248 | pairID:124, seed_elecID:22, subject:8 | 1.14 | 11.47 | <<0.01 | <<0.01 |
| midTH | OCC | 32 | pairID:16, seed_elecID:5, subject:3 | 0.64 | 2.53 | 0.02 | 0.03 |
| midTH | antTH | 42 | pairID:21, seed_elecID:16, subject:10 | 0.24 | 0.93 | 0.34 | 0.39 |
| midTH | midTH | 26 | pairID:13, seed_elecID:8, subject:6 | -0.18 | -0.67 | 0.49 | 0.53 |
| midTH | pstTH | 74 | pairID:37, seed_elecID:24, subject:10 | 0.2 | 1.2 | 0.23 | 0.29 |
| pstTH | BG | 334 | pairID:166, seed_elecID:61, subject:16 | 0.11 | 1.2 | 0.23 | 0.29 |
| pstTH | AMY | 52 | pairID:26, seed_elecID:17, subject:8 | 0.68 | 2.91 | <<0.01 | 0.01 |
| pstTH | OFC | 94 | pairID:47, seed_elecID:15, subject:9 | 0.6 | 3.1 | <<0.01 | 0.01 |
| pstTH | ACC | 64 | pairID:32, seed_elecID:8, subject:5 | 1.64 | 7.56 | <<0.01 | <<0.01 |
| pstTH | MFC | 50 | pairID:25, seed_elecID:10, subject:5 | 0.98 | 5.1 | <<0.01 | <<0.01 |
| pstTH | LFC | 570 | pairID:280, seed_elecID:55, subject:17 | 0.43 | 6.46 | <<0.01 | <<0.01 |
| pstTH | INS | 526 | pairID:261, seed_elecID:54, subject:19 | 0.14 | 1.83 | 0.07 | 0.11 |
| pstTH | TP | 20 | pairID:10, seed_elecID:5, subject:3 | 1.16 | 2.27 | 0.03 | 0.05 |
| pstTH | HPC | 114 | pairID:56, seed_elecID:24, subject:16 | -0.21 | -1.26 | 0.21 | 0.28 |
| pstTH | PHG | 26 | pairID:13, seed_elecID:7, subject:5 | 0.77 | 3.42 | <<0.01 | 0.01 |
| pstTH | FG | 62 | pairID:31, seed_elecID:9, subject:5 | 0.17 | 1.02 | 0.3 | 0.35 |
| pstTH | ITG | 138 | pairID:68, seed_elecID:21, subject:11 | 0.15 | 1.15 | 0.25 | 0.31 |
| pstTH | MTG | 354 | pairID:176, seed_elecID:39, subject:16 | -0.05 | -0.58 | 0.56 | 0.58 |
| pstTH | STG | 392 | pairID:193, seed_elecID:41, subject:13 | -0.73 | -7.72 | <<0.01 | <<0.01 |
| pstTH | MCC | 72 | pairID:36, seed_elecID:14, subject:5 | 0.59 | 3.95 | <<0.01 | <<0.01 |
| pstTH | SM | 462 | pairID:231, seed_elecID:38, subject:11 | 0.36 | 4.64 | <<0.01 | <<0.01 |
| *pstTH* | SPL | 64 | pairID:29, seed_elecID:10, subject:5 | -0.32 | -1.81 | 0.07 | 0.11 |
| *pstTH* | IPL | 312 | pairID:154, seed_elecID:37, subject:13 | -0.12 | -1.37 | 0.17 | 0.25 |
| *pstTH* | PMC | 588 | pairID:293, seed_elecID:43, subject:15 | -0.08 | -1.34 | 0.18 | 0.25 |
| *pstTH* | OCC | 76 | pairID:38, seed_elecID:10, subject:5 | 0.52 | 2.66 | 0.01 | 0.02 |
| *pstTH* | antTH | 120 | pairID:60, seed_elecID:36, subject:17 | 0.11 | 1.04 | 0.3 | 0.35 |
| *pstTH* | midTH | 92 | pairID:46, seed_elecID:24, subject:12 | 0.1 | 0.66 | 0.51 | 0.55 |
| *pstTH* | pstTH | 46 | pairID:23, seed_elecID:18, subject:10 | 0.12 | 0.63 | 0.52 | 0.55 |

<<: far smaller than; If the Estimate value is above zero, it means F3-outflow > F1-outflow

**Table S7.** Significance testing results for the comparison of the late thalamocortical pathway (F3-outflow) > early corticothalamic (F1-inflow) pathway in different brain areas of both hemispheres. The tests used a mixed linear model, with the z-scored r* as the dependent variable, predicated by (i) the fixed effect of *pathway* (i.e., “F3-outflow” vs. “F1-inflow”), (ii) the random effect of the coupled-contacts identity for a connectivity pair (i.e., the repeated measure of the electrode contacts concerned in both inflow and outflow pathways), (iii) the random effect of the thalamic sites being considered (ii) the random effect of individual differences. Model comparison is conducted to test the significance of the fixed effect of *pathway*. Using the toolbox of “lme4” ^56^ (https://cran.r-project.org/web/packages/lme4/), the model and the compared null model are written as the following: ml: zr ∼ pathway + (1 | pairID) + (1 | seed_elecID) + (1 | subject); ml0: zr ∼ (1 | pairID) + (1 | seed_elecID) + (1 | subject).

| ROI | Area | NO.<br>Observations | NO.<br>Random Groups (adjusted) | Estimate | t-value | p | p (FDR-corr.) |
| --- | --- | --- | --- | --- | --- | --- | --- |
| antTH | BG | 261 | pairID:210, seed_elecID:65, subject:19 | 0.01 | 0.05 | 0.96 | 0.98 |
| antTH | AMY | 113 | pairID:102, seed_elecID:50, subject:17 | 0.03 | 0.18 | 0.87 | 0.93 |
| antTH | OFC | 390 | pairID:332, seed_elecID:59, subject:19 | 0.28 | 3.12 | 0.00 | 0.01 |
| antTH | ACC | 200 | pairID:167, seed_elecID:36, subject:10 | -0.14 | -1.17 | 0.25 | 0.39 |
| antTH | MFC | 67 | pairID:58, seed_elecID:21, subject:8 | 0.07 | 0.42 | 0.58 | 0.70 |
| antTH | LFC | 900 | pairID:786, seed_elecID:76, subject:23 | 0.23 | 3.57 | 0.00 | 0.00 |
| antTH | INS | 519 | pairID:446, seed_elecID:79, subject:24 | 0.09 | 1.09 | 0.27 | 0.40 |
| antTH | TP | 85 | NA | NA | NA | NA | NA |
| antTH | HPC | 206 | pairID:190, seed_elecID:60, subject:21 | -0.30 | -2.51 | 0.02 | 0.04 |
| antTH | PHG | 33 | pairID:28, seed_elecID:17, subject:8 | 0.01 | 0.04 | 0.98 | 0.98 |
| antTH | FG | 64 | pairID:59, seed_elecID:29, subject:11 | 0.00 | 0.00 | 0.98 | 0.98 |
| antTH | ITG | 132 | pairID:122, seed_elecID:41, subject:17 | 0.75 | 4.88 | <<0.01 | <<0.01 |
| antTH | MTG | 346 | pairID:316, seed_elecID:55, subject:21 | 0.59 | 6.13 | <<0.01 | <<0.01 |
| antTH | STG | 188 | pairID:173, seed_elecID:47, subject:18 | -0.28 | -1.81 | 0.07 | 0.15 |
| antTH | MCC | 76 | pairID:58, seed_elecID:19, subject:6 | -0.25 | -1.61 | 0.14 | 0.25 |
| antTH | SM | 201 | pairID:186, seed_elecID:39, subject:14 | 0.42 | 2.54 | 0.01 | 0.03 |
| antTH | SPL | 39 | NA | NA | NA | NA | NA |
| antTH | IPL | 141 | pairID:126, seed_elecID:36, subject:12 | 0.77 | 4.62 | <<0.01 | <<0.01 |
| antTH | PMC | 333 | pairID:304, seed_elecID:61, subject:18 | 0.55 | 6.16 | <<0.01 | <<0.01 |
| antTH | OCC | 58 | NA | NA | NA | NA | NA |
| antTH | antTH | 18 | NA | NA | NA | NA | NA |
| antTH | midTH | 45 | pairID:41, seed_elecID:24, subject:11 | -0.15 | -0.91 | 0.42 | 0.57 |
| antTH | pstTH | 139 | pairID:116, seed_elecID:54, subject:18 | 0.17 | 1.29 | 0.20 | 0.32 |
| midTH | BG | 174 | pairID:143, seed_elecID:38, subject:10 | -0.10 | -0.79 | 0.43 | 0.58 |
| midTH | AMY | 72 | pairID:66, seed_elecID:33, subject:9 | 0.09 | 0.42 | 0.65 | 0.74 |
| midTH | OFC | 244 | pairID:217, seed_elecID:41, subject:10 | 0.11 | 1.01 | 0.33 | 0.48 |
| midTH | ACC | 140 | pairID:118, seed_elecID:24, subject:7 | 0.50 | 4.55 | <<0.01 | <<0.01 |
| midTH | MFC | 87 | pairID:80, seed_elecID:27, subject:7 | 0.61 | 3.75 | 0.00 | 0.00 |
| midTH | LFC | 590 | pairID:518, seed_elecID:49, subject:12 | 0.56 | 8.01 | <<0.01 | <<0.01 |
| midTH | INS | 286 | pairID:253, seed_elecID:50, subject:13 | 0.34 | 3.86 | 0.00 | 0.00 |
| midTH | TP | 30 | NA | NA | NA | NA | NA |
| midTH | HPC | 100 | pairID:96, seed_elecID:44, subject:12 | -0.28 | -1.09 | 0.27 | 0.40 |
| midTH | PHG | 10 | NA | NA | NA | NA | NA |
| midTH | FG | 16 | NA | NA | NA | NA | NA |
| midTH | ITG | 69 | pairID:64, seed_elecID:24, subject:7 | 1.08 | 4.49 | <<0.01 | <<0.01 |
| midTH | MTG | 162 | pairID:149, seed_elecID:36, subject:11 | 0.68 | 4.52 | <<0.01 | <<0.01 |
| midTH | STG | 67 | pairID:63, seed_elecID:30, subject:11 | -0.11 | -0.58 | 0.55 | 0.67 |
| midTH | MCC | 63 | pairID:53, seed_elecID:22, subject:6 | -0.09 | -0.46 | 0.61 | 0.71 |
| midTH | SM | 238 | pairID:211, seed_elecID:33, subject:9 | 0.60 | 4.83 | <<0.01 | <<0.01 |
| midTH | SPL | 27 | NA | NA | NA | NA | NA |
| midTH | IPL | 94 | NA | NA | NA | NA | NA |
| midTH | PMC | 197 | pairID:187, seed_elecID:44, subject:10 | 0.48 | 3.00 | 0.00 | 0.01 |
| midTH | OCC | 21 | NA | NA | NA | NA | NA |
| midTH | antTH | 45 | pairID:41, seed_elecID:30, subject:11 | -0.14 | -0.77 | 0.49 | 0.64 |
| midTH | midTH | 26 | NA | NA | NA | NA | NA |
| midTH | pstTH | 83 | pairID:73, seed_elecID:39, subject:12 | 0.03 | 0.20 | 0.84 | 0.92 |
| pstTH | BG | 284 | pairID:250, seed_elecID:95, subject:16 | 0.08 | 0.64 | 0.52 | 0.66 |
| pstTH | AMY | 110 | pairID:102, seed_elecID:54, subject:13 | -0.32 | -1.32 | 0.20 | 0.32 |
| pstTH | OFC | 157 | pairID:147, seed_elecID:55, subject:14 | 0.67 | 4.12 | <<0.01 | 0.00 |
| pstTH | ACC | 87 | pairID:78, seed_elecID:26, subject:8 | 0.90 | 4.33 | <<0.01 | 0.00 |
| pstTH | MFC | 70 | NA | NA | NA | NA | NA |
| pstTH | LFC | 604 | pairID:547, seed_elecID:100, subject:19 | 0.36 | 4.54 | <<0.01 | <<0.01 |
| pstTH | INS | 570 | pairID:497, seed_elecID:102, subject:20 | 0.18 | 1.99 | 0.05 | 0.10 |
| pstTH | TP | 53 | NA | NA | NA | NA | NA |
| pstTH | HPC | 209 | pairID:194, seed_elecID:75, subject:18 | -0.62 | -4.72 | <<0.01 | <<0.01 |
| pstTH | PHG | 45 | pairID:39, seed_elecID:20, subject:6 | 0.14 | 0.68 | 0.53 | 0.66 |
| pstTH | FG | 64 | pairID:59, seed_elecID:27, subject:9 | -0.10 | -0.38 | 0.68 | 0.77 |
| pstTH | ITG | 170 | pairID:156, seed_elecID:47, subject:13 | 0.42 | 2.38 | 0.02 | 0.05 |
| pstTH | MTG | 341 | pairID:311, seed_elecID:66, subject:17 | -0.19 | -1.80 | 0.08 | 0.16 |
| pstTH | STG | 319 | pairID:290, seed_elecID:75, subject:15 | -0.21 | -1.63 | 0.11 | 0.20 |
| pstTH | MCC | 85 | pairID:69, seed_elecID:27, subject:6 | 0.22 | 1.30 | 0.19 | 0.32 |
| pstTH | SM | 426 | pairID:356, seed_elecID:54, subject:12 | 0.25 | 2.97 | <<0.01 | 0.01 |
| pstTH | SPL | 66 | pairID:58, seed_elecID:23, subject:7 | 0.02 | 0.10 | 0.88 | 0.93 |
| pstTH | IPL | 258 | pairID:225, seed_elecID:66, subject:13 | 0.22 | 1.90 | 0.06 | 0.13 |
| *pstTH* | PMC | 556 | pairID:475, seed_elecID:78, subject:15 | 0.24 | 3.07 | <<0.01 | 0.01 |
| *pstTH* | OCC | 106 | pairID:99, seed_elecID:27, subject:5 | -0.33 | -1.24 | 0.34 | 0.48 |
| *pstTH* | antTH | 139 | pairID:116, seed_elecID:71, subject:18 | 0.22 | 1.73 | 0.11 | 0.20 |
| *pstTH* | midTH | 83 | pairID:73, seed_elecID:39, subject:12 | 0.35 | 2.34 | 0.02 | 0.05 |
| *pstTH* | *pstTH* | 46 | NA | NA | NA | NA | NA |

<<: far smaller than; If the Estimate value is above zero, it means F3-outflow >F1-inflow

## REFERENCES

1 Sporns, O., Chialvo, D. R., Kaiser, M. & Hilgetag, C. C. Organization, development and function of complex brain networks. Trends Cogn. Sci. 8, 418–425 (2004).

2 Shine, J. M., Lewis, L. D., Garrett, D. D. & Hwang, K. The impact of the human thalamus on brain-wide information processing. Nature reviews. Neuroscience (2023). 10.1038/s41583-023-00701-0

3 Matsumoto, R. et al. Functional connectivity in the human language system: a cortico-cortical evoked potential study. Brain 127, 2316–2330 (2004).

4 Matsumoto, R. et al. Functional connectivity in human cortical motor system: a cortico-cortical evoked potential study. Brain 130, 181–197 (2007).

5 Wu, T. Q. et al. Multisite thalamic recordings to characterize seizure propagation in the human brain. Brain 146, 2792–2802 (2023). 10.1093/brain/awad121

6 Zumsteg, D., Lozano, A. M. & Wennberg, R. A. Depth electrode recorded cerebral responses with deep brain stimulation of the anterior thalamus for epilepsy. Clin Neurophysiol 117, 1602–1609 (2006). 10.1016/j.clinph.2006.04.008

7 Ojeda Valencia, G., et al. Signatures of Electrical Stimulation Driven Network Interactions in the Human Limbic System. The Journal of Neuroscience: The Official Journal of the Society for Neuroscience 43, 6697–6711 (2023). 10.1523/JNEUROSCI.2201-22.2023

8 Miller, K. J., Muller, K. R. & Hermes, D. Basis profile curve identification to understand electrical stimulation effects in human brain networks. PLoS Comput Biol 17, e1008710 (2021). 10.1371/journal.pcbi.1008710

9 Jedynak, M. et al. Variability of Single Pulse Electrical Stimulation Responses Recorded with Intracranial Electroencephalography in Epileptic Patients. Brain Topogr 36, 119–127 (2023). 10.1007/s10548-022-00928-7

10 Becht, E. et al. Dimensionality reduction for visualizing single-cell data using UMAP. Nat Biotechnol (2018). 10.1038/nbt.4314

11 Stieger, J. R., et al. Cross regional coordination of neural activity in the human brain during autobiographical self-referential processing. PNAS In Press (2023).

12 Kunieda, T., Yamao, Y., Kikuchi, T. & Matsumoto, R. New Approach for Exploring Cerebral Functional Connectivity: Review of Cortico-cortical Evoked Potential. Neurologia medico-chirurgica 55, 374–382 (2015). 10.2176/nmc.ra.2014-0388

13 Keller, C. J. et al. Intrinsic functional architecture predicts electrically evoked responses in the human brain. Proceedings of the National Academy of Sciences 108, 10308–10313 (2011). 10.1073/pnas.1019750108

14 Keller, C. J. et al. Mapping human brain networks with cortico-cortical evoked potentials. Philos Trans R Soc Lond B Biol Sci 369, 20130528 (2014). 10.1098/rstb.2013.0528

15 Togo, M. et al. Distinct connectivity patterns in human medial parietal cortices: Evidence from standardized connectivity map using cortico-cortical evoked potential. NeuroImage 263, 119639 (2022). 10.1016/j.neuroimage.2022.119639

16 Fogerson, P. M. & Huguenard, J. R. Tapping the Brakes: Cellular and Synaptic Mechanisms that Regulate Thalamic Oscillations. Neuron 92, 687–704 (2016). 10.1016/j.neuron.2016.10.024

17 Buzsaki, G. Rhythms of the Brain. (Oxford University Press, 2006).

18 Groppe, D. M. et al. Dominant frequencies of resting human brain activity as measured by the electrocorticogram. Neuroimage 79, 223–233 (2013). 10.1016/j.neuroimage.2013.04.044

19 Hacker, C. D., Snyder, A. Z., Pahwa, M., Corbetta, M. & Leuthardt, E. C. Frequency-specific electrophysiologic correlates of resting state fMRI networks. Neuroimage 149, 446–457 (2017). 10.1016/j.neuroimage.2017.01.054

20 Fisher, R. et al. Electrical stimulation of the anterior nucleus of thalamus for treatment of refractory epilepsy. Epilepsia 51, 899–908 (2010). 10.1111/j.1528-1167.2010.02536.x

21 Engel, J., Jr. & Pitkanen, A. Biomarkers for epileptogenesis and its treatment. Neuropharmacology 167, 107735 (2020). 10.1016/j.neuropharm.2019.107735

22 Caciagli, L., Bernhardt, B. C., Hong, S.-J., Bernasconi, A. & Bernasconi, N. Functional network alterations and their structural substrate in drug-resistant epilepsy. Front Neurosci 8, 411 (2014). 10.3389/fnins.2014.00411

23 Fleury, M. et al. Episodic memory network connectivity in temporal lobe epilepsy. Epilepsia 63, 2597–2622 (2022). 10.1111/epi.17370

24 Pittau, F., Grova, C., Moeller, F., Dubeau, F. & Gotman, J. Patterns of altered functional connectivity in mesial temporal lobe epilepsy. Epilepsia 53, 1013–1023 (2012). 10.1111/j.1528-1167.2012.03464.x

25 Roger, E. et al. Hubs disruption in mesial temporal lobe epilepsy. A resting-state fMRI study on a language-and-memory network. Human Brain Mapping 41, 779–796 (2020). 10.1002/hbm.24839

26 Li, L. et al. Topographical reorganization of brain functional connectivity during an early period of epileptogenesis. Epilepsia 62, 1231–1243 (2021). 10.1111/epi.16863

27 Mazrooyisebdani, M. et al. Graph Theory Analysis of Functional Connectivity Combined with Machine Learning Approaches Demonstrates Widespread Network Differences and Predicts Clinical Variables in Temporal Lobe Epilepsy. Brain Connect 10, 39–50 (2020). 10.1089/brain.2019.0702

28 Ofer, I. et al. Association between seizure freedom and default mode network reorganization in patients with unilateral temporal lobe epilepsy. Epilepsy Behav 90, 238–246 (2019). 10.1016/j.yebeh.2018.10.025

29 Liao, W. et al. Default mode network abnormalities in mesial temporal lobe epilepsy: A study combining fMRI and DTI. Human Brain Mapping 32, 883–895 (2011). 10.1002/hbm.21076

30 Widjaja, E., Zamyadi, M., Raybaud, C., Snead, O. C. & Smith, M. L. Abnormal Functional Network Connectivity among Resting-State Networks in Children with Frontal Lobe Epilepsy. AJNR Am J Neuroradiol 34, 2386–2392 (2013). 10.3174/ajnr.A3608

31 Zhang, Z. et al. Impaired attention network in temporal lobe epilepsy: A resting FMRI study. Neuroscience Letters 458, 97–101 (2009). 10.1016/j.neulet.2009.04.040

32 Keller, C. J. et al. Mapping human brain networks with cortico-cortical evoked potentials. Philos Trans R Soc Lond B Biol Sci 369 (2014). 10.1098/rstb.2013.0528

33 Veit, M. J. et al. Temporal order of signal propagation within and across intrinsic brain networks. Proc Natl Acad Sci U S A 118 (2021). 10.1073/pnas.2105031118

34 Seguin, C. et al. Communication dynamics in the human connectome shape the cortex-wide propagation of direct electrical stimulation. Neuron 0 (2023). 10.1016/j.neuron.2023.01.027

35 Guo, Z. H. et al. Epileptogenic network of focal epilepsies mapped with cortico-cortical evoked potentials. Clin Neurophysiol 131, 2657–2666 (2020). 10.1016/j.clinph.2020.08.012

36 Arthuis, M. et al. Impaired consciousness during temporal lobe seizures is related to increased long-distance cortical-subcortical synchronization. Brain 132, 2091–2101 (2009). 10.1093/brain/awp086

37 Guye, M. et al. The role of corticothalamic coupling in human temporal lobe epilepsy. Brain 129, 1917–1928 (2006). 10.1093/brain/awl151

38 Pizzo, F. et al. The Ictal Signature of Thalamus and Basal Ganglia in Focal Epilepsy: A SEEG Study. Neurology 96, e280–e293 (2021). 10.1212/WNL.0000000000011003

39 Filipescu, C. et al. The effect of medial pulvinar stimulation on temporal lobe seizures. Epilepsia 60, e25–e30 (2019). 10.1111/epi.14677

40 Evangelista, E. et al. Does the Thalamo-Cortical Synchrony Play a Role in Seizure Termination? Front Neurol 6, 192 (2015). 10.3389/fneur.2015.00192

41 Gadot, R., Korst, G., Shofty, B., Gavvala, J. R. & Sheth, S. A. Thalamic stereoelectroencephalography in epilepsy surgery: a scoping literature review. J Neurosurg, 1–16 (2022). 10.3171/2022.1.JNS212613

42 Ilyas, A., Tandon, N. & Lhatoo, S. D. Thalamic neuromodulation for epilepsy: A clinical perspective. Epilepsy Res 183, 106942 (2022). 10.1016/j.eplepsyres.2022.106942

43 Chaitanya, G. et al. Robot-assisted stereoelectroencephalography exploration of the limbic thalamus in human focal epilepsy: implantation technique and complications in the first 24 patients. Neurosurg Focus 48, E2 (2020). 10.3171/2020.1.FOCUS19887

44 Romeo, A. et al. Early ictal recruitment of midline thalamus in mesial temporal lobe epilepsy. Ann Clin Transl Neurol 6, 1552–1558 (2019). 10.1002/acn3.50835

45 McKhann, G. M. Editorial. Dulling the double-edged sword of human SEEG research. Neurosurg Focus 48, E3 (2020). 10.3171/2020.1.FOCUS2069

46 Groppe, D. M. et al. iELVis: An open source MATLAB toolbox for localizing and visualizing human intracranial electrode data. J Neurosci Methods 281, 40–48 (2017). 10.1016/j.jneumeth.2017.01.022

47 Fischl, B. FreeSurfer. NeuroImage 62, 774–781 (2012).

48 Jenkinson, M., Beckmann, C. F., Behrens, T. E., Woolrich, M. W. & Smith, S. M. Fsl. Neuroimage 62, 782–790 (2012). 10.1016/j.neuroimage.2011.09.015

49 Jenkinson, M. & Smith, S. A global optimisation method for robust affine registration of brain images. Medical image analysis 5, 143–156 (2001).

50 Greve, D. N. & Fischl, B. Accurate and robust brain image alignment using boundary- based registration. Neuroimage 48, 63–72 (2009). 10.1016/j.neuroimage.2009.06.060

51 Papademetris, X. et al. BioImage Suite: An integrated medical image analysis suite: An update. The insight journal 2006, 209 (2006).

52 Su, J. H. et al. Thalamus Optimized Multi Atlas Segmentation (THOMAS): fast, fully automated segmentation of thalamic nuclei from structural MRI. Neuroimage 194, 272–282 (2019). 10.1016/j.neuroimage.2019.03.021

53 Liu, S. & Parvizi, J. Cognitive refractory state caused by spontaneous epileptic high-frequency oscillations in the human brain. Sci Transl Med 11 (2019). 10.1126/scitranslmed.aax7830

54 McInnes, L., Healy, J. & Melville, J. (arXiv, 2020).

55 Campello, R. J. G. B., Moulavi, D. & Sander, J. (eds Jian Pei et al.) 160–172 (Springer).

56 Bates, D., Mächler, M., Bolker, B. & Walker, S. Fitting Linear Mixed-Effects Models Using lme4. Journal of Statistical Software 67, 1–48 (2015). 10.18637/jss.v067.i01

